## Supplementary Information for "Human class B1 GPCR modulation by plasma membrane lipids"

Kin W. Chao *et al.*

**This file includes:**

Figs. S1 to S20
Tables S1, S2

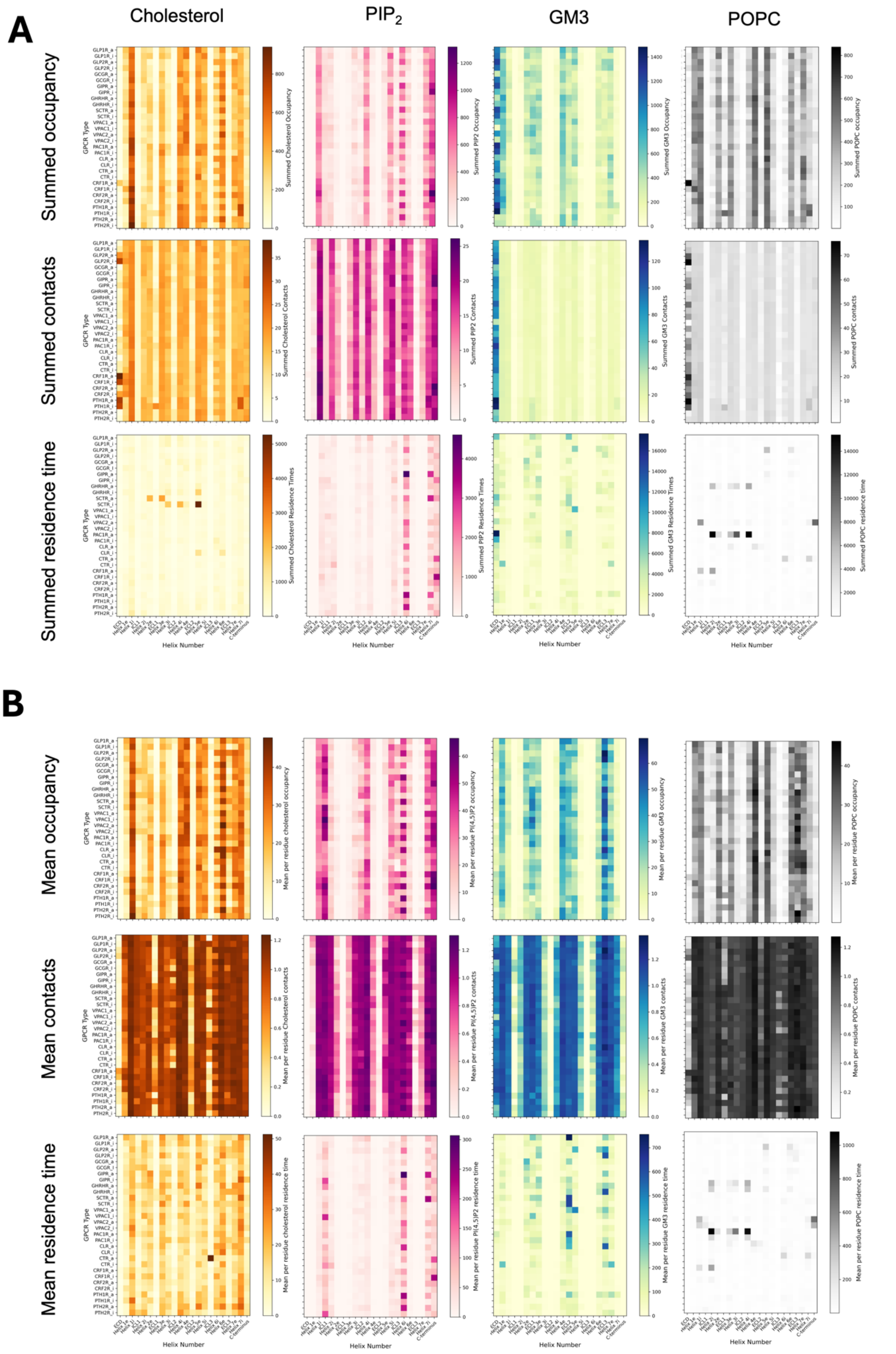
A
Cholesterol
$PIP_2$
GM3
POPC
Summed occupancy
Summed contacts
Summed residence time
B
Mean occupancy
Mean contacts
Mean residence time
GPCR Type
Helix Number

**Fig. S1.**

**<u>Global analysis of class B1 GPCR interactions with the regulatory lipids cholesterol, $PIP_2$ and GM3</u>, and presumed non-regulatory lipid POPC.** Lipid occupancy (**A**), contacts with membrane (B) and lipid residence time (**C**) is shown for each class B1 GPCR as a function of ECD, helix, and loop number. These properties were computed over 3 x 10 μs of production run simulation time using PyLipID [31] and summed (top) or averaged (lower) over the constituent residues of each region. Helix 1e denotes the extracellular half of Helix 1, and Helix 1i the intracellular half. Active states are denoted by _a, and inactive by _i. The residue ranges defining each region correspond to those from GPCRdb [32].

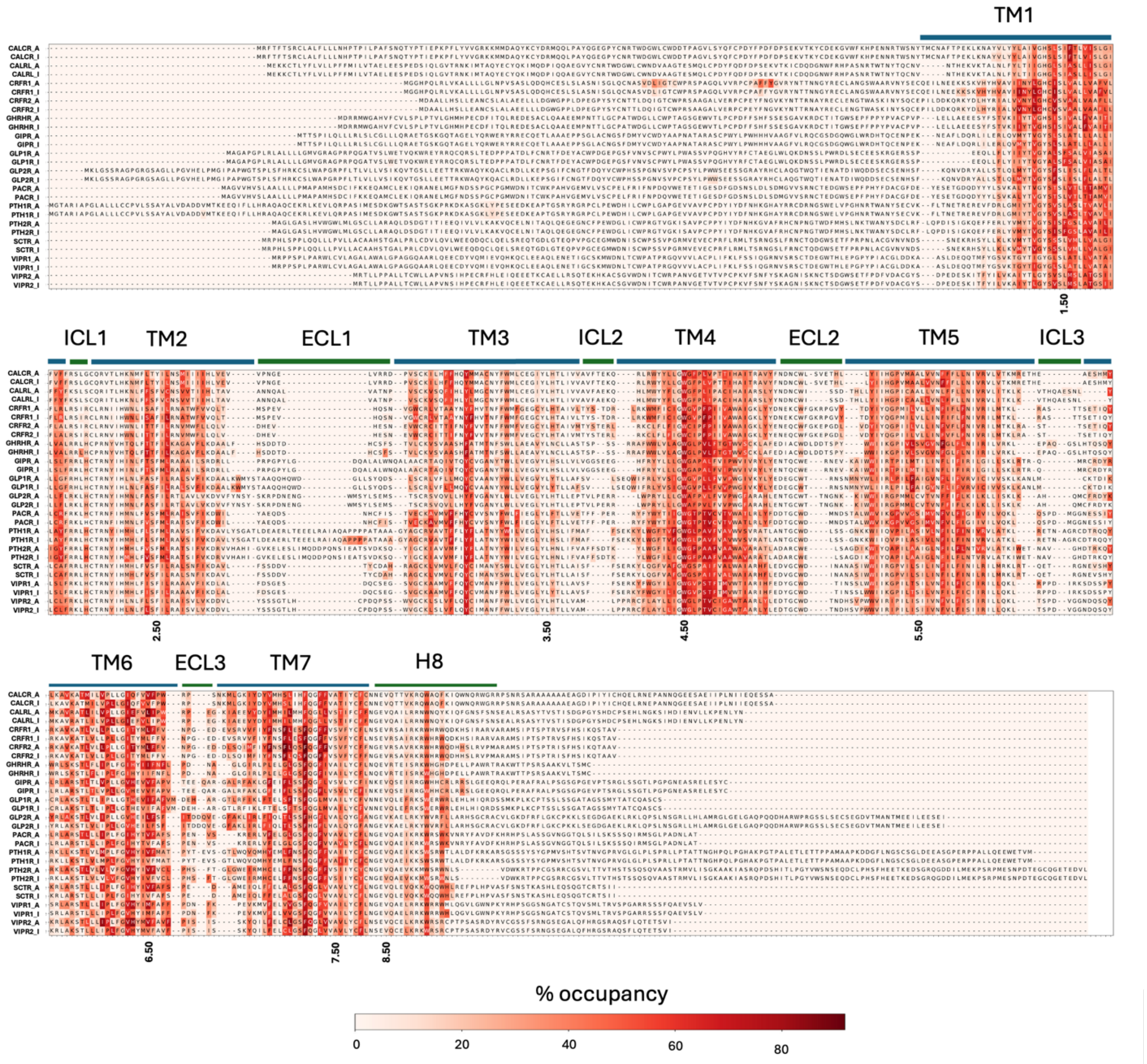


**Fig. S2.**

**Cholesterol interactions with class B1 GPCRs.** The calculated cholesterol occupancy values from cgMD simulations (averaged across three repeats, using a cutoff = 0.7 nm) for active and inactive states are mapped onto the aligned sequence and shown as a heatmap for cholesterol for the 15 receptors in the B1 family (high occupancy to low occupancy: red to white.

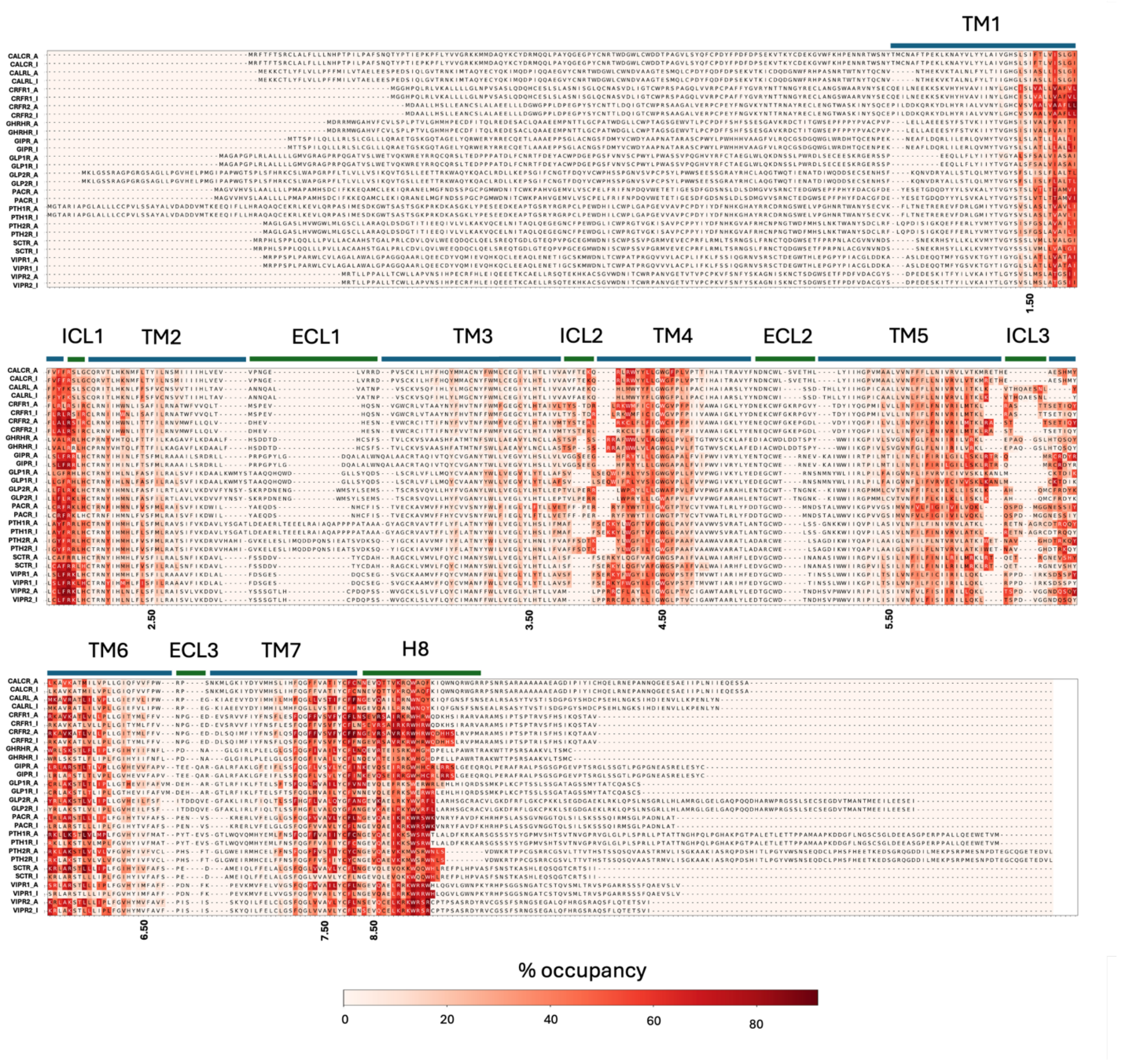


**Fig. S3.**

**PIP2 interactions with class B1 GPCRs.** The calculated PIP2 occupancy values from cgMD simulations (averaged across three repeats, using a cutoff = 0.7 nm) for active and inactive states are mapped onto the aligned sequence and shown as a heatmap for PIP2 for the 15 receptors in the B1 family (high occupancy to low occupancy: red to white.

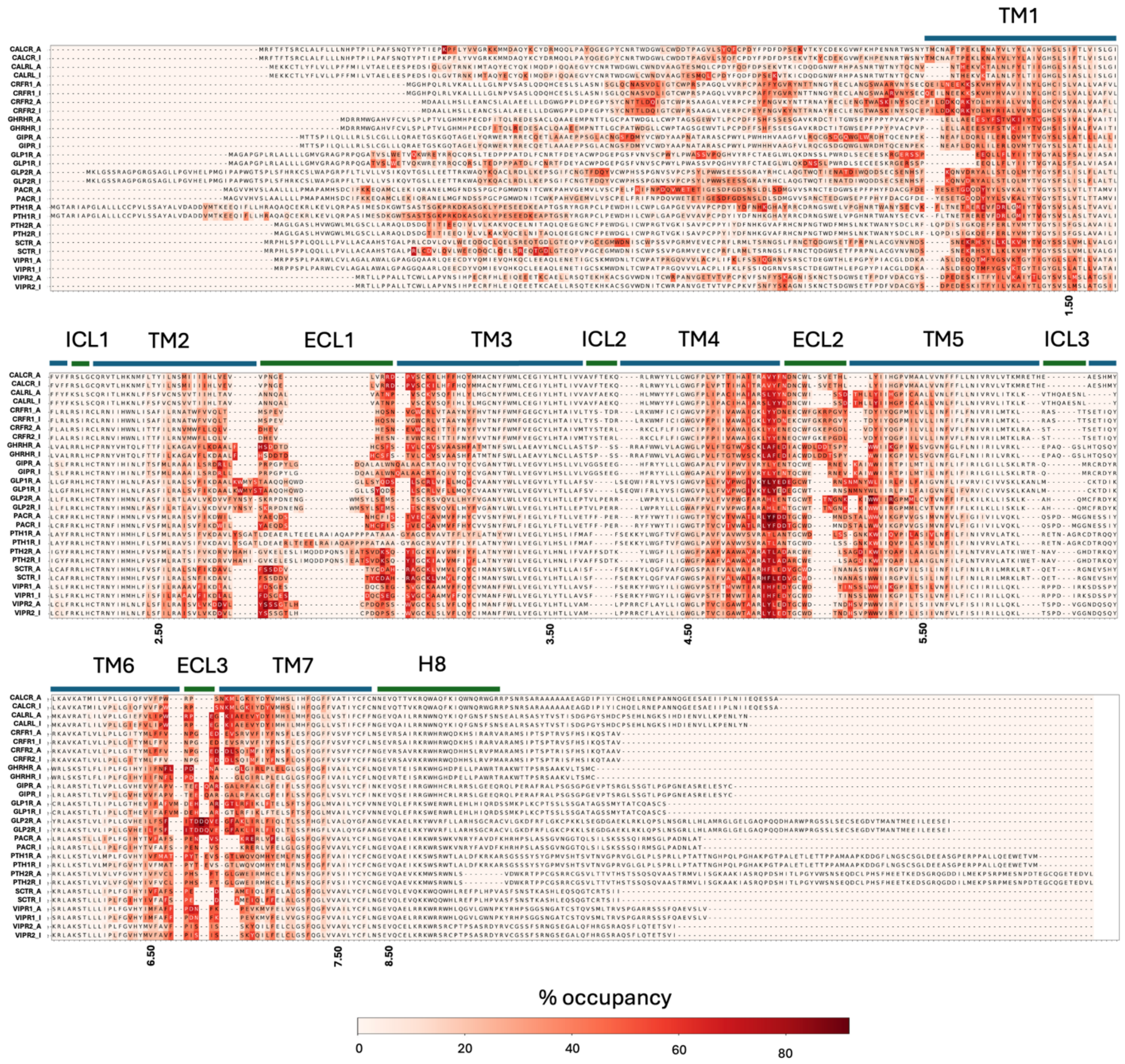


**Fig. S4.**

**GM3 interactions with class B1 GPCRs.** The calculated cholesterol occupancy values from cgMD simulations (averaged across three repeats, using a cutoff = 0.7 nm) for active and inactive states are mapped onto the aligned sequence and shown as a heatmap for GM3 for the 15 receptors in the B1 family (high occupancy to low occupancy: red to white).

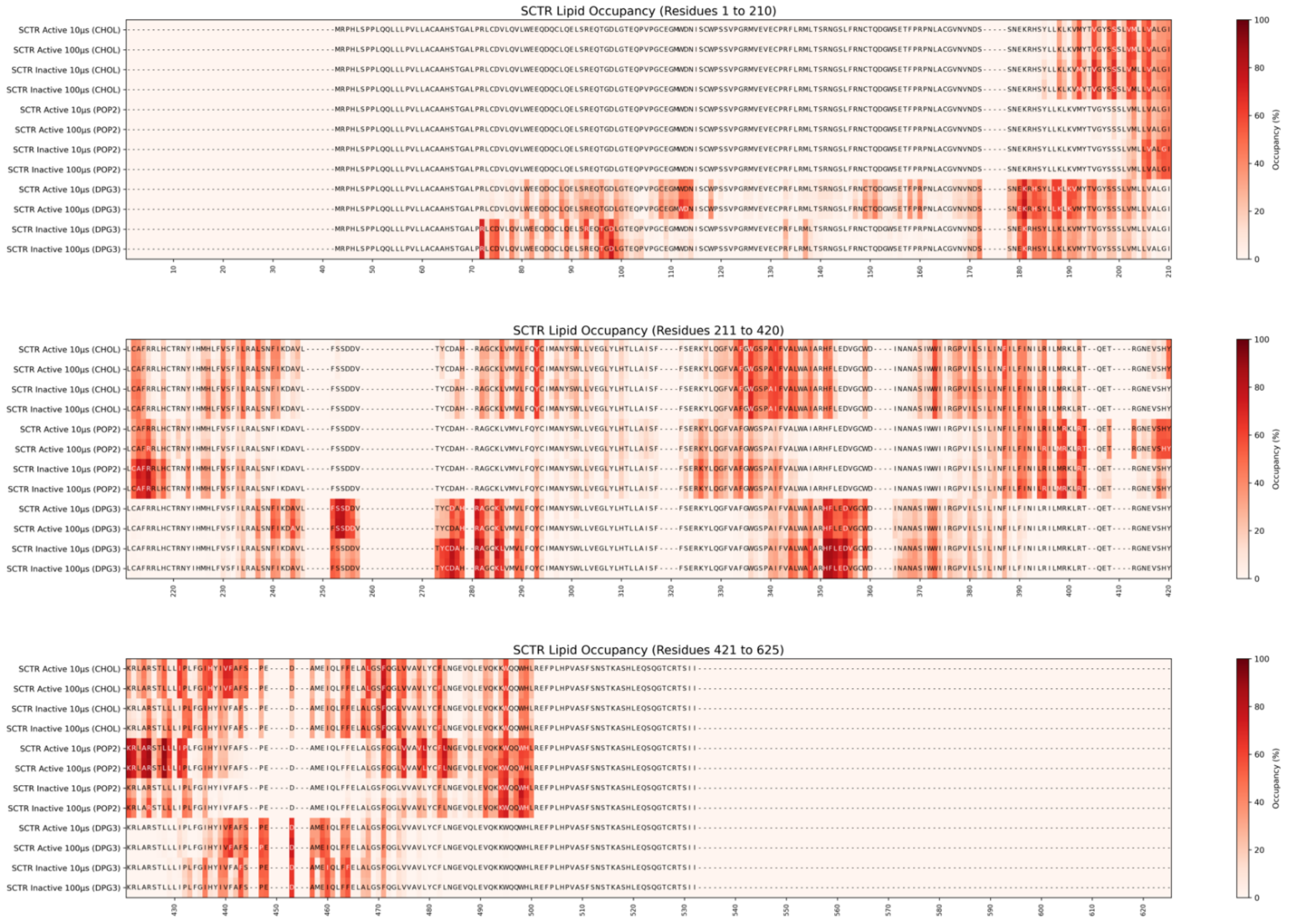


**Fig. S5.**

**Cholesterol, PIP$_2$ and GM3 occupancy for SCTR in 10 µs vs 100 µs simulations.** The calculated lipid occupancy values from cgMD simulations (averaged across three repeats, using a cutoff = 0.7 nm) for active and inactive states are mapped onto the aligned sequence and shown as a heatmap for GM3 for SCTR in the active and inactive states. The 100 µs simulations were generated from independent membrane configurations. High occupancy to low occupancy: red to white. Martini3 naming: POP2 = PIP$_2$, DPG3 = GM3.

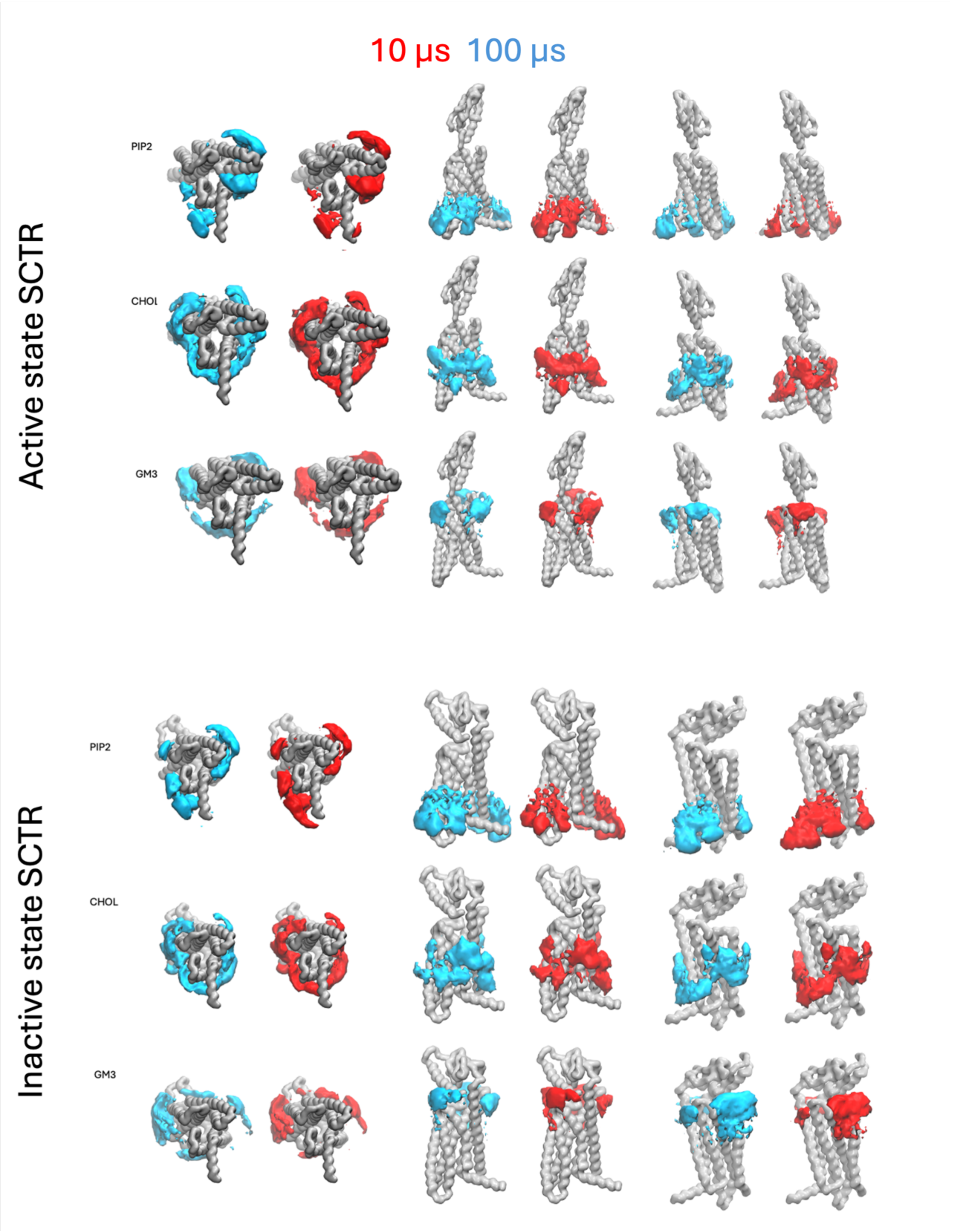


**Fig. S6.**

**Cholesterol, $PIP_2$ and GM3 volumetric density maps showing lipid headgroup positions relative to SCTR in 10 µs vs 100 µs simulations.** Maps were generated from three combined trajectories. Volumes are shown in blue (100 µs) and red (10 µs).

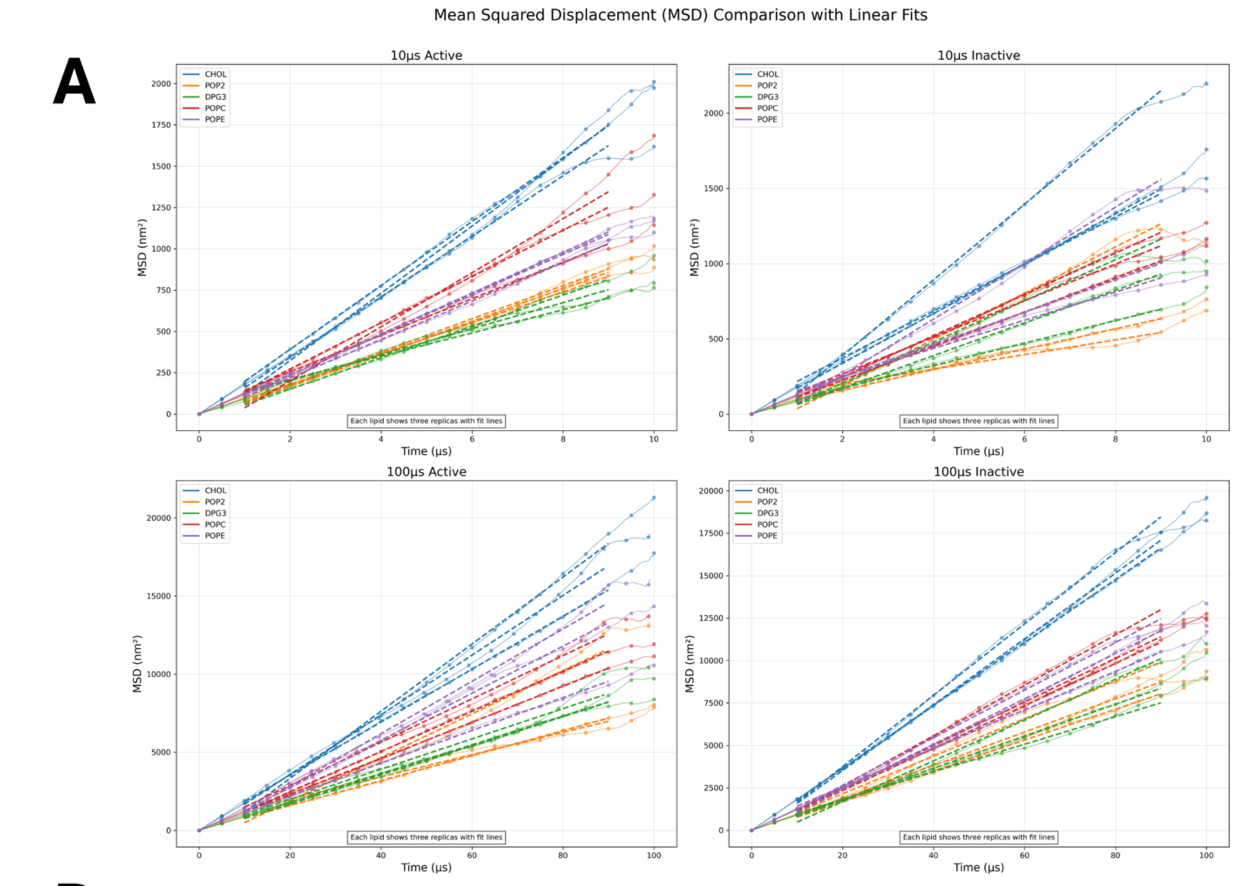

Mean Squared Displacement (MSD) Comparison with Linear Fits
A
10µs Active
10µs Inactive
100µs Active
100µs Inactive
CHOL
POP2
DPG3
POPC
POPE
MSD (nm²)
Time (µs)
Each lipid shows three replicas with fit lines

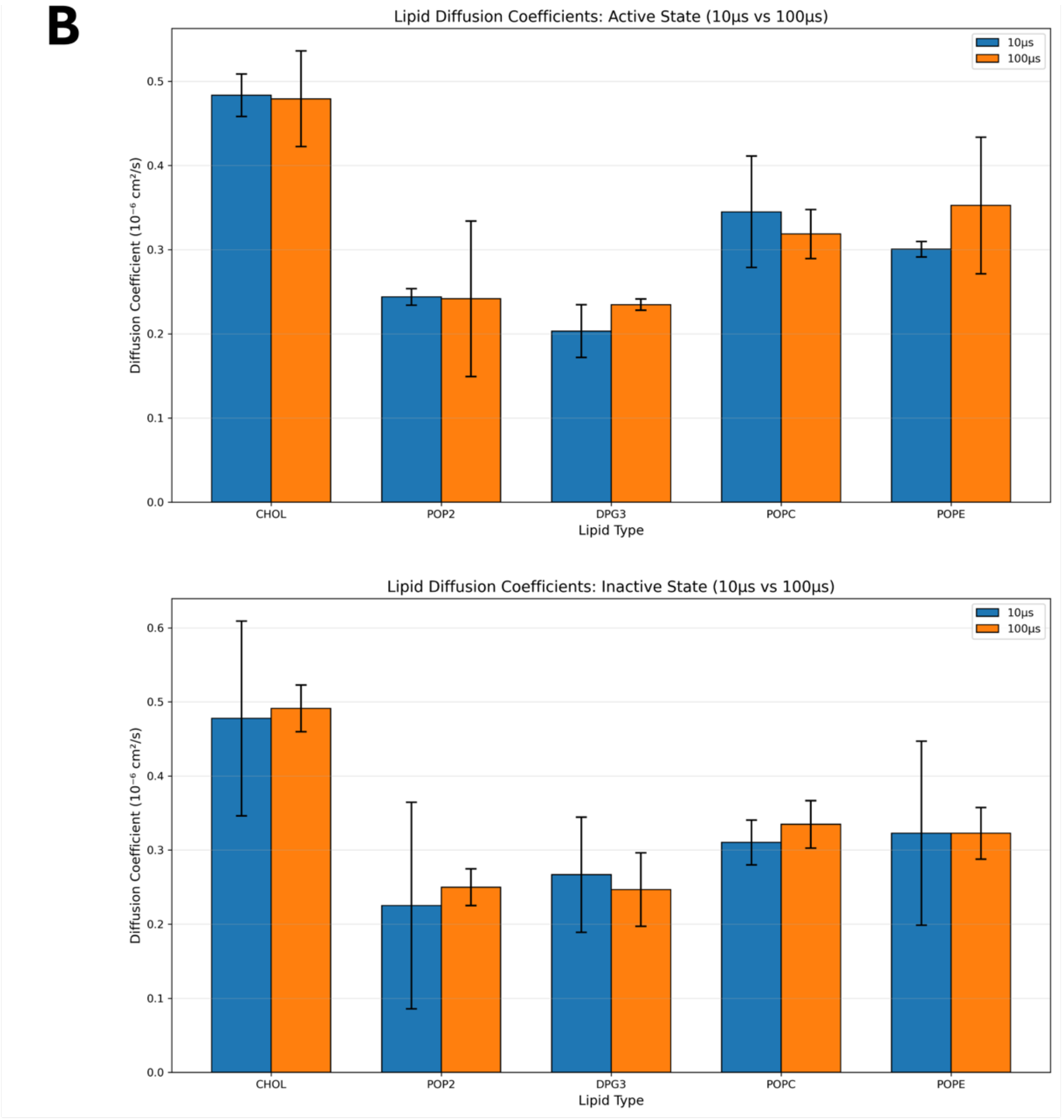


**Fig. S7. Lipid diffusion properties**. **A)** MSD against time calculated using gmx msd for the SCTR active (left) and inactive (right) states for 10 μs (top) vs 100 μs (bottom). The calculation was performed using all frames in the simulations, and the raw data points are marked at regular intervals. A linear line was fitted to the 10-90% data range, which was used to obtain the diffusion coefficients. **B)** Diffusion coefficients for different lipid types (CHOL, $PIP_2$, GM3, POPC, and POPE) comparing 10 μs (blue) vs 100 μs (orange) simulations in the SCTR active (top) and inactive (bottom) states. The diffusion coefficients were obtained from the linear fit of MSD data shown in Figure X. Error bars represent standard deviation across three independent replicas. The diffusion coefficients are presented in units of $10^{-6}$ cm²/s. Martini3 naming: POP2 = $PIP_2$, DPG3 = GM3

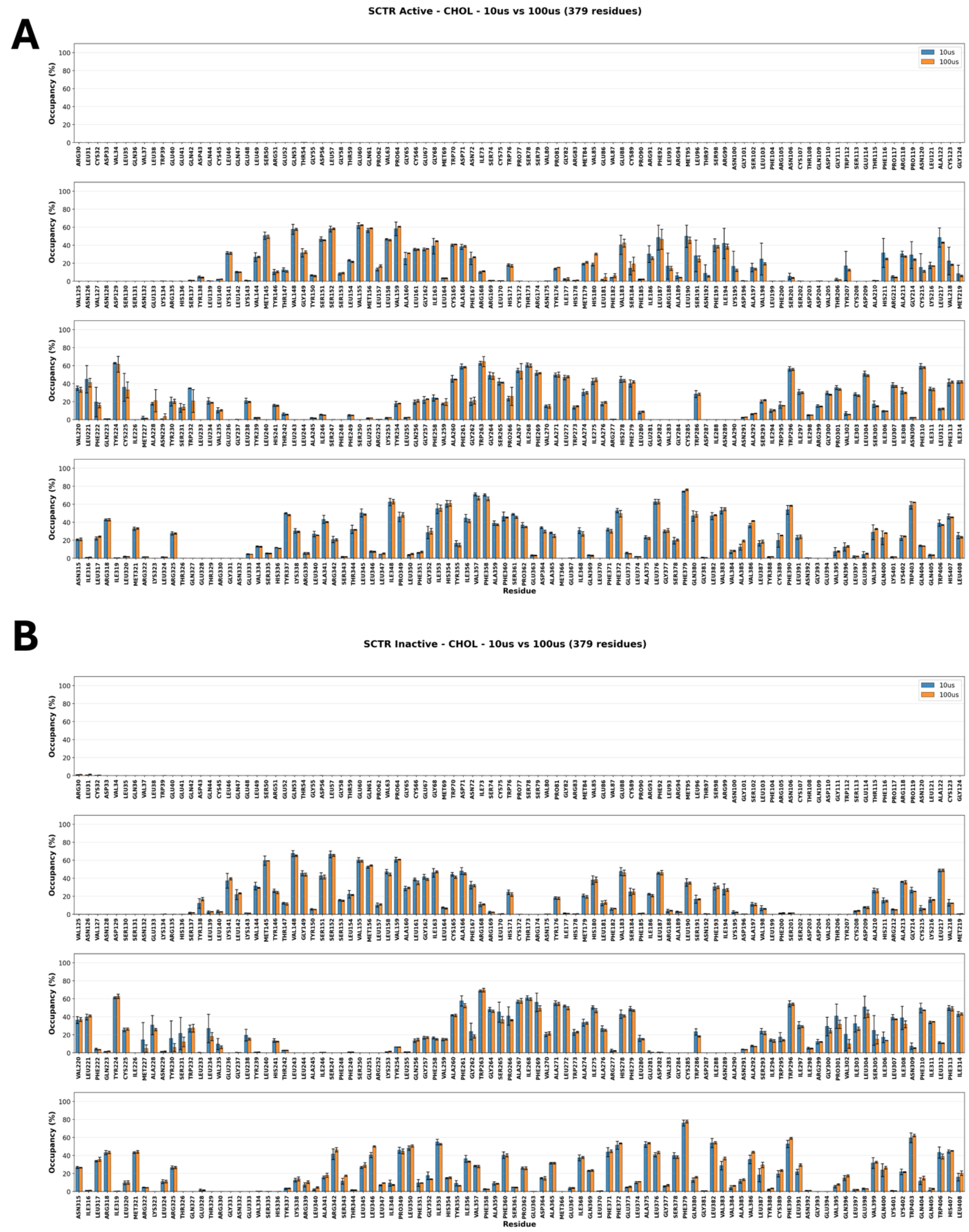


**Fig. S8. Cholesterol occupancy error estimates for SCTR**. Bar chart showing mean lipid occupancy per residue for active and inactive states of SCTR calculated from three independent simulation repeats (R1, R2, R3) using 7.0 Å cutoff distance for cholesterol. Error bars represent standard deviation across the three repeats.

CLR_active
Cholesterol

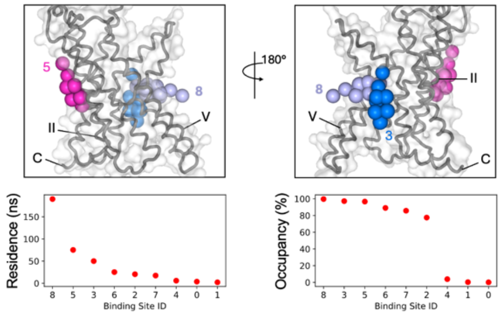


CLR_inactive
Cholesterol

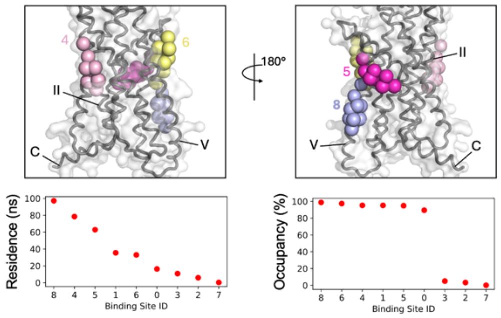


CLR_active
PIP2

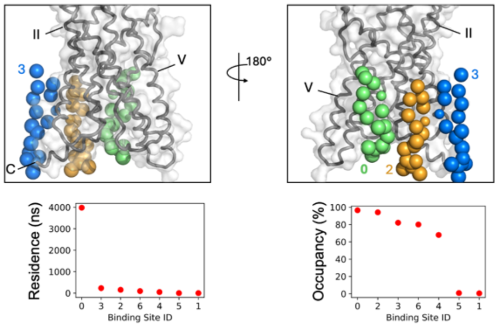


CLR_inactive
PIP2

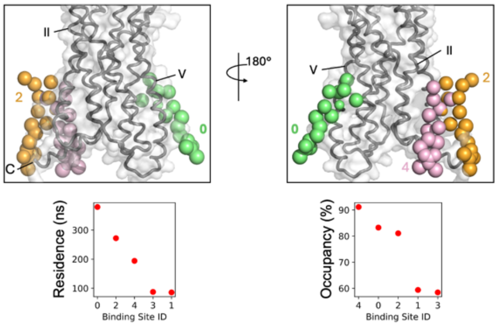


CRF1R_active
Cholesterol

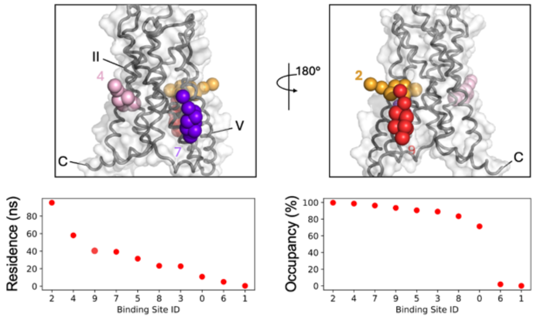


CRF1R_inactive
Cholesterol

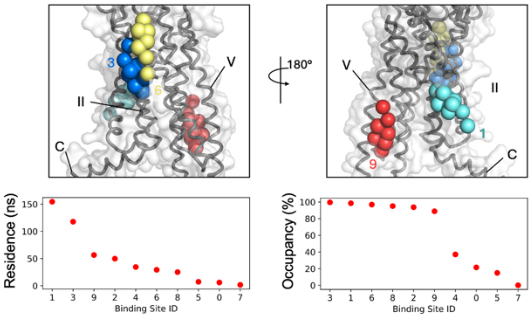


CRF1R_active
PIP2

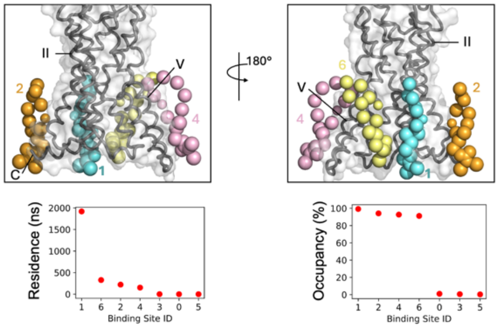


CRF1R_inactive
PIP2

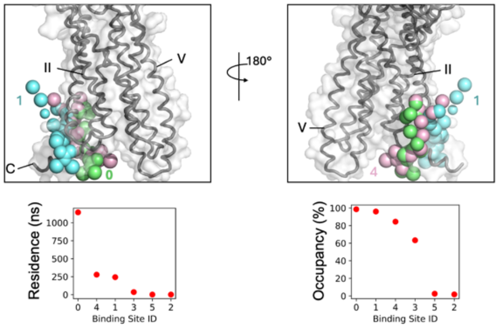

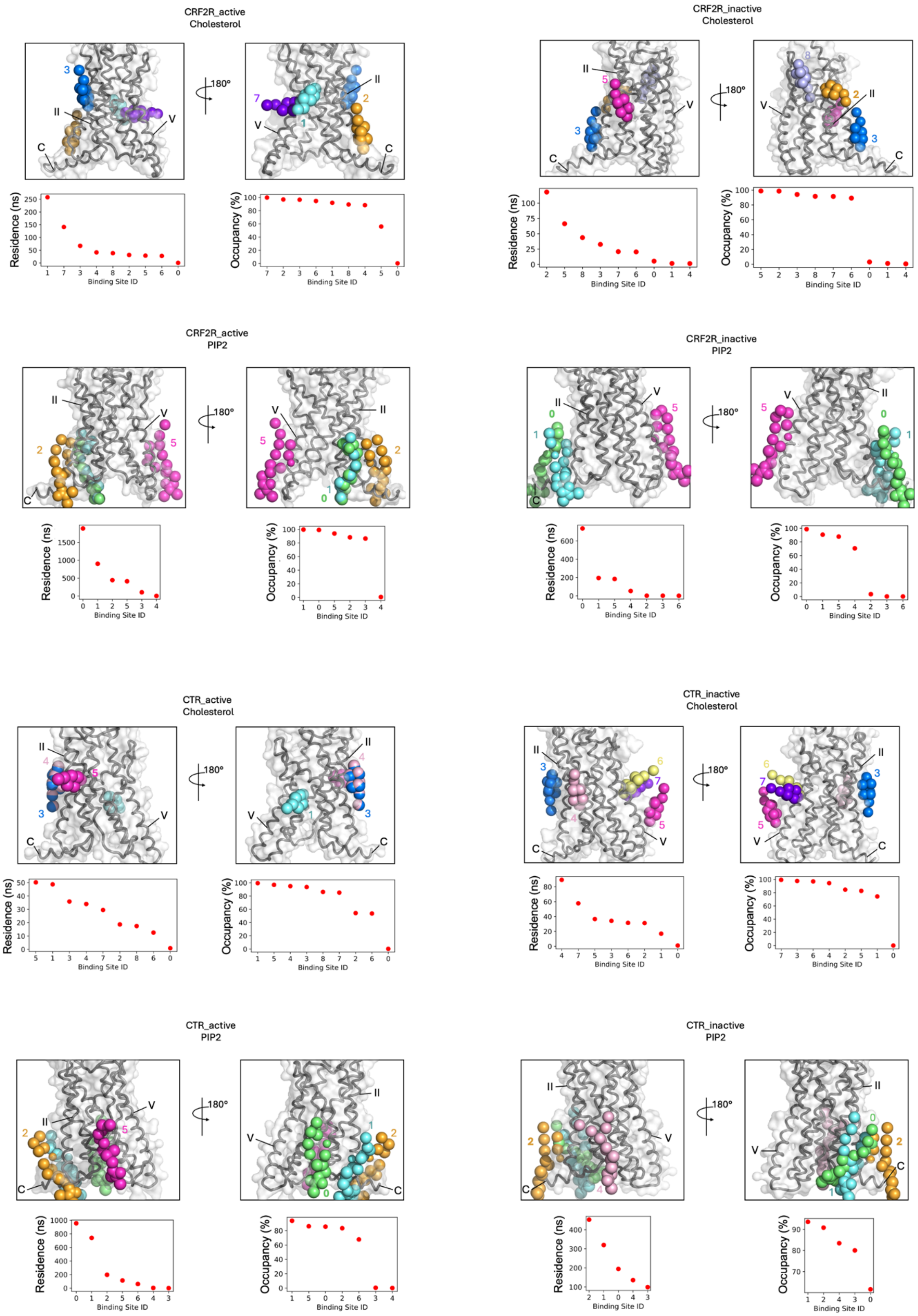
CRF2R_active
Cholesterol
180°
Residence (ns)
Occupancy (%)
Binding Site ID
CRF2R_inactive
Cholesterol
CRF2R_active
PIP2
CRF2R_inactive
PIP2
CTR_active
Cholesterol
CTR_inactive
Cholesterol
CTR_active
PIP2
CTR_inactive
PIP2

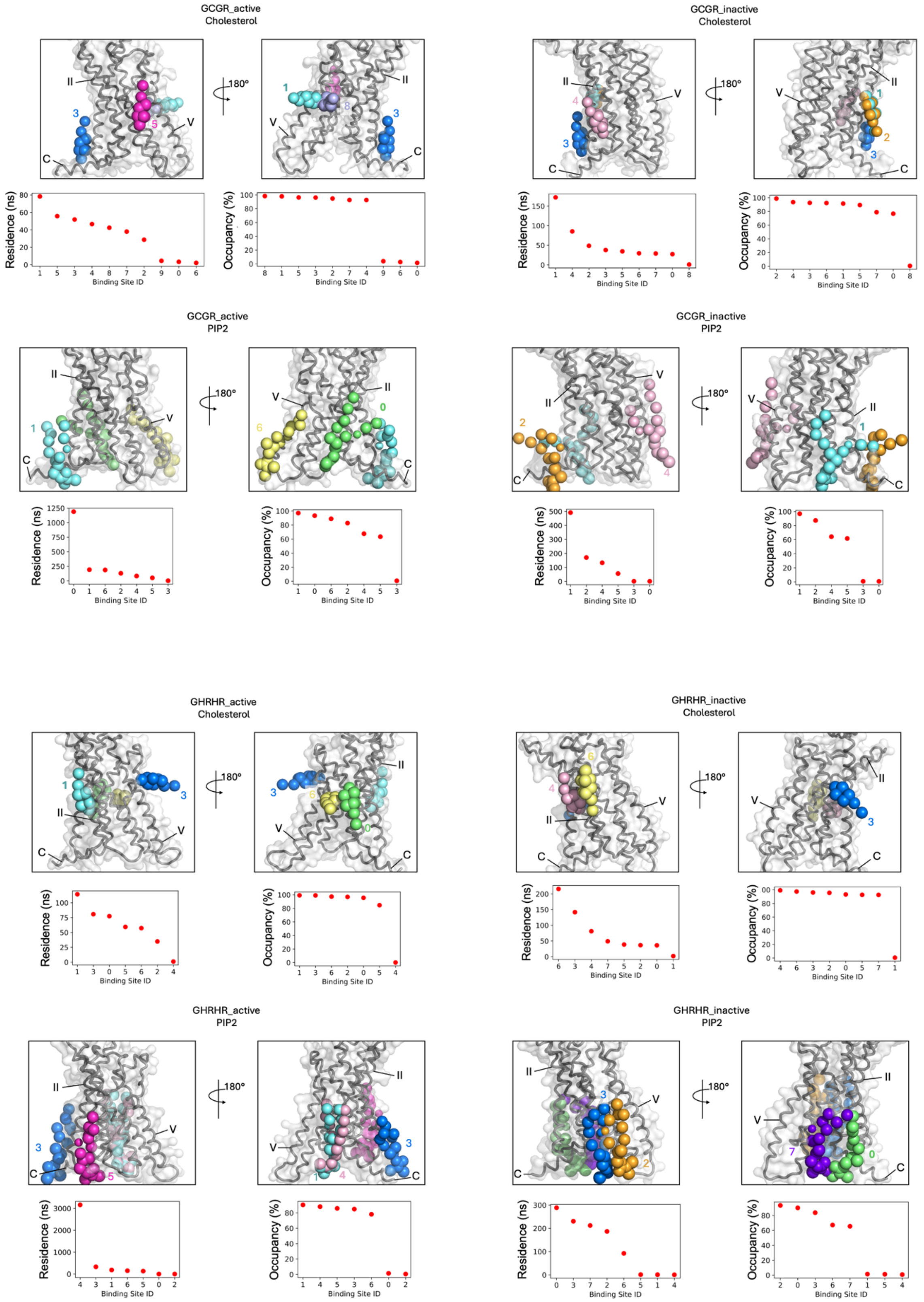
GCGR_active
Cholesterol
180°
Residence (ns)
Occupancy (%)
Binding Site ID
GCGR_inactive
Cholesterol
GCGR_active
PIP2
GCGR_inactive
PIP2
GHRHR_active
Cholesterol
GHRHR_inactive
Cholesterol
GHRHR_active
PIP2
GHRHR_inactive
PIP2

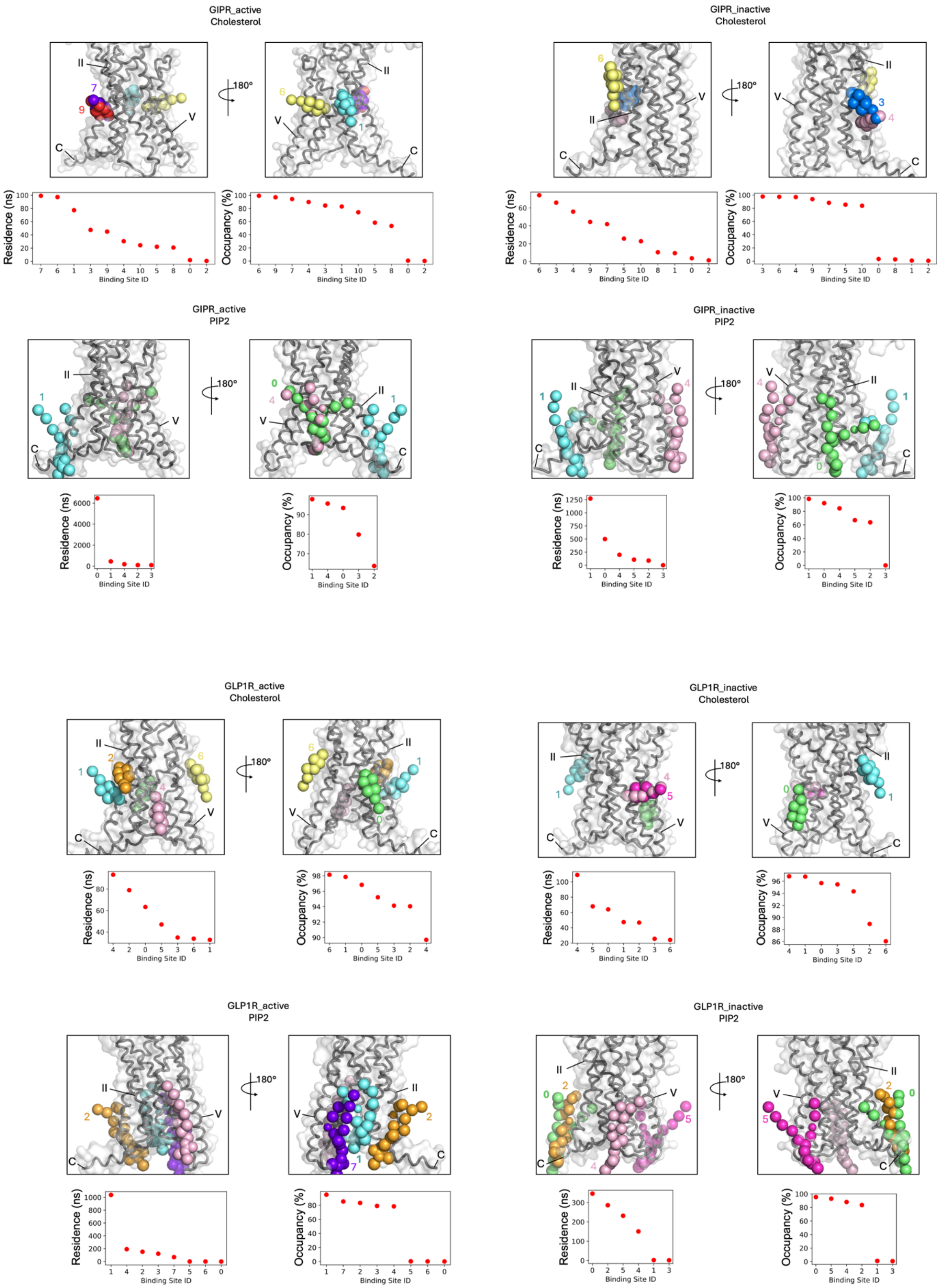
GIPR_active
Cholesterol
GIPR_inactive
Cholesterol
GIPR_active
PIP2
GIPR_inactive
PIP2
GLP1R_active
Cholesterol
GLP1R_inactive
Cholesterol
GLP1R_active
PIP2
GLP1R_inactive
PIP2
180°
Residence (ns)
Occupancy (%)
Binding Site ID

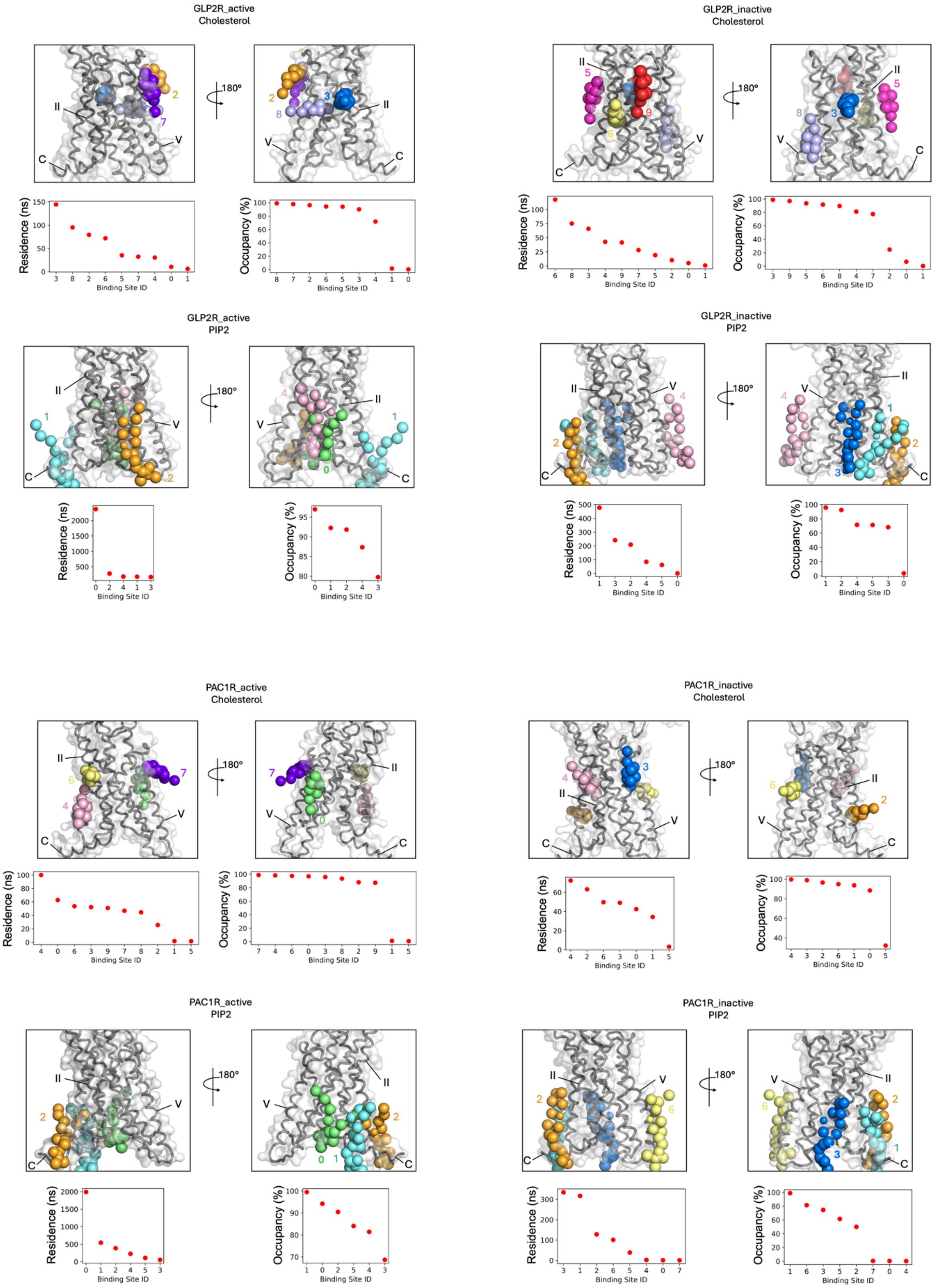
GLP2R_active
Cholesterol
GLP2R_inactive
Cholesterol
GLP2R_active
PIP2
GLP2R_inactive
PIP2
PAC1R_active
Cholesterol
PAC1R_inactive
Cholesterol
PAC1R_active
PIP2
PAC1R_inactive
PIP2
180°
Residence (ns)
Occupancy (%)
Binding Site ID

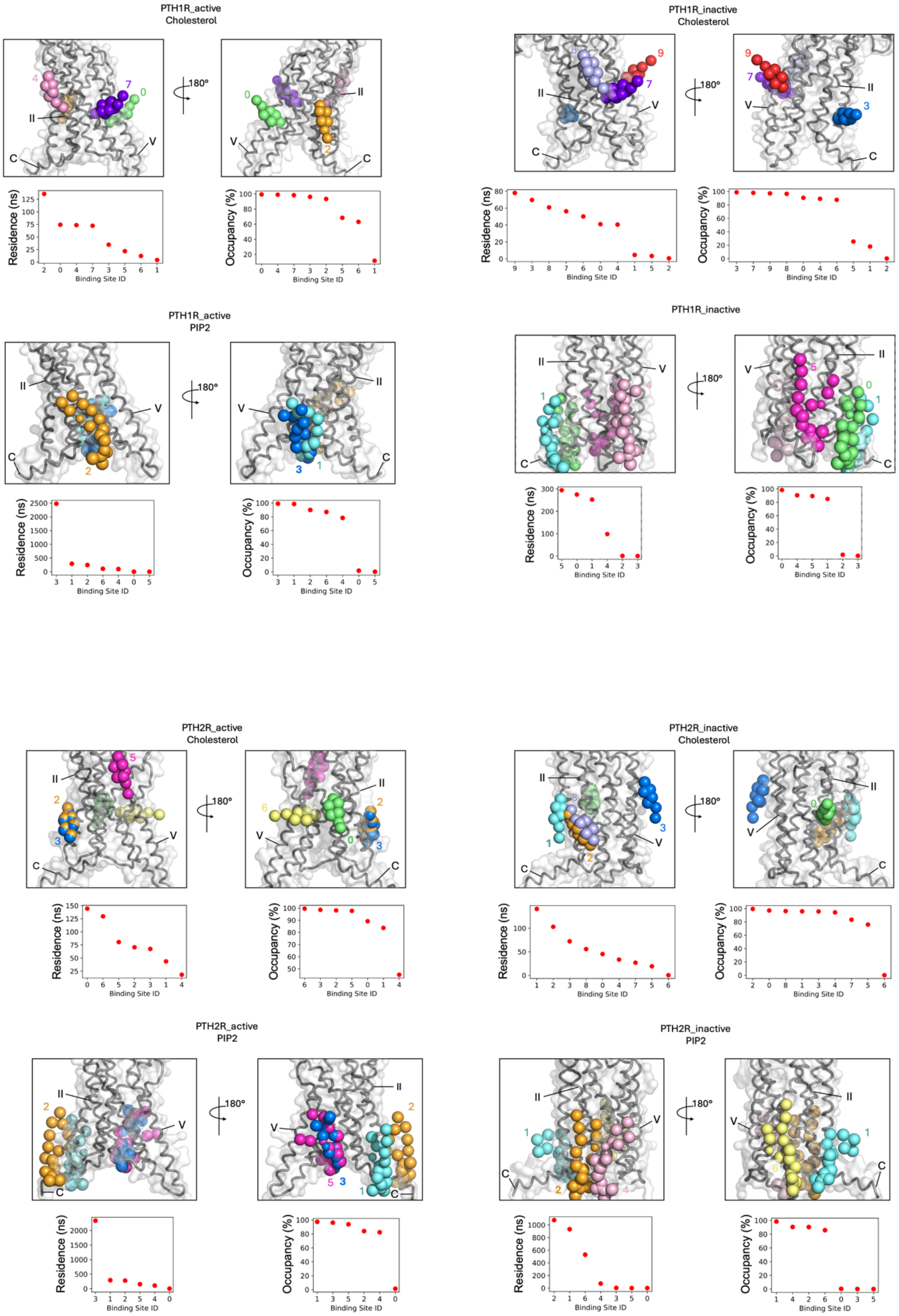
PTH1R_active
Cholesterol
PTH1R_inactive
Cholesterol
PTH1R_active
PIP2
PTH1R_inactive
PTH2R_active
Cholesterol
PTH2R_inactive
Cholesterol
PTH2R_active
PIP2
PTH2R_inactive
PIP2
180°
Residence (ns)
Occupancy (%)
Binding Site ID

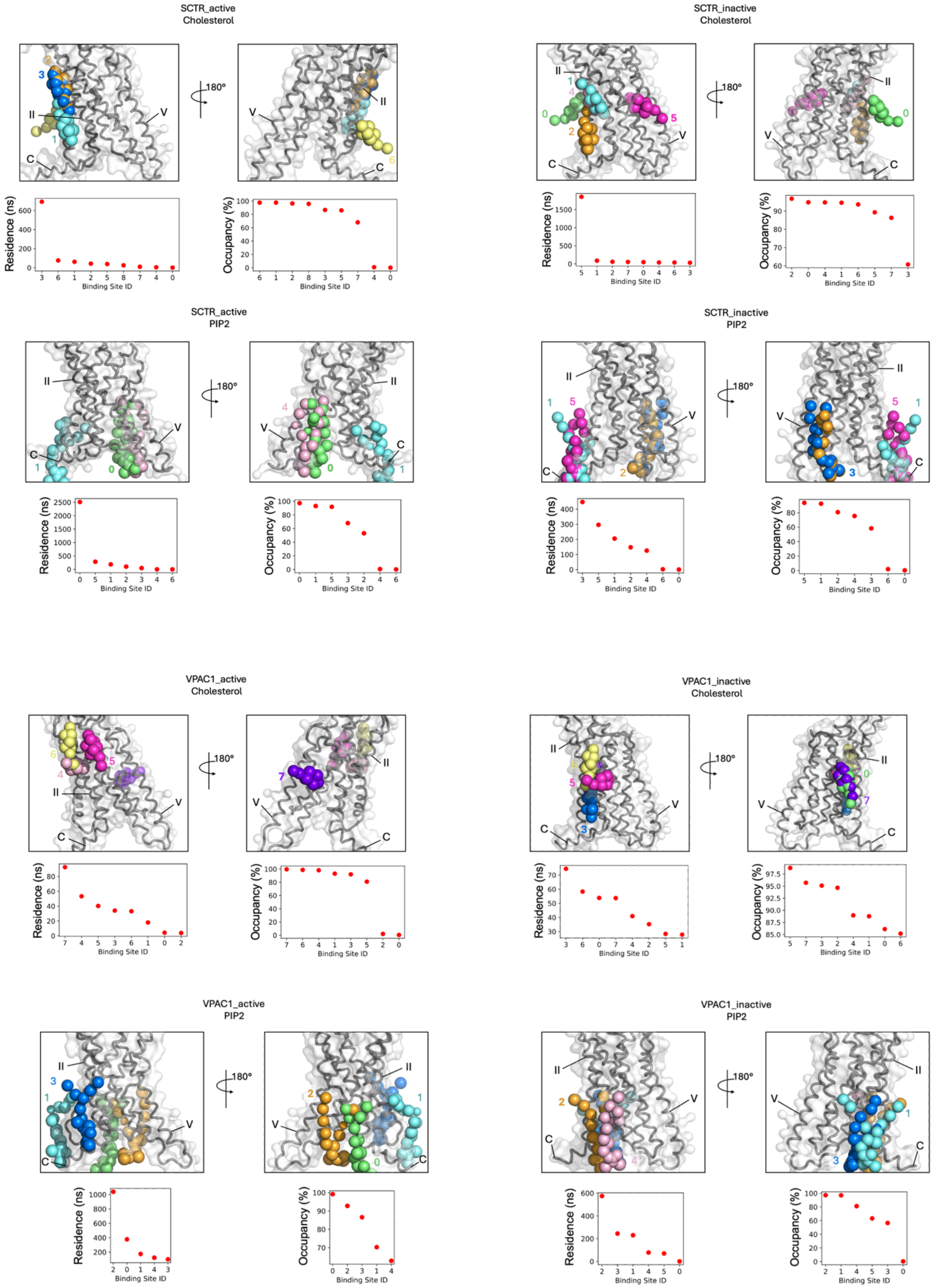

SCTR_active
Cholesterol
180°
Residence (ns)
Occupancy (%)
Binding Site ID
SCTR_inactive
Cholesterol
SCTR_active
PIP2
SCTR_inactive
PIP2
VPAC1_active
Cholesterol
VPAC1_inactive
Cholesterol
VPAC1_active
PIP2
VPAC1_inactive
PIP2

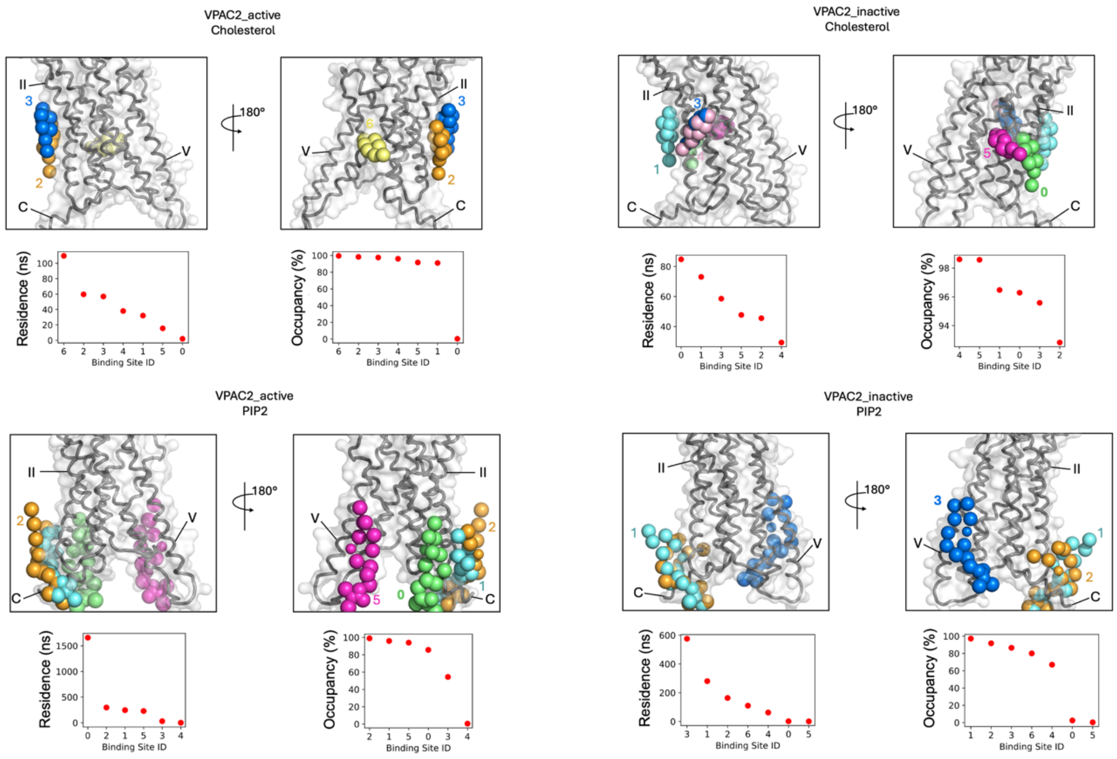


**Fig. S9.**

**Cholesterol and $PIP_2$ binding sites for class B1 receptors**. The top 3 lipid binding poses by residence time and occupancy are shown and labelled according to site ID in the residence time and occupancy plots.

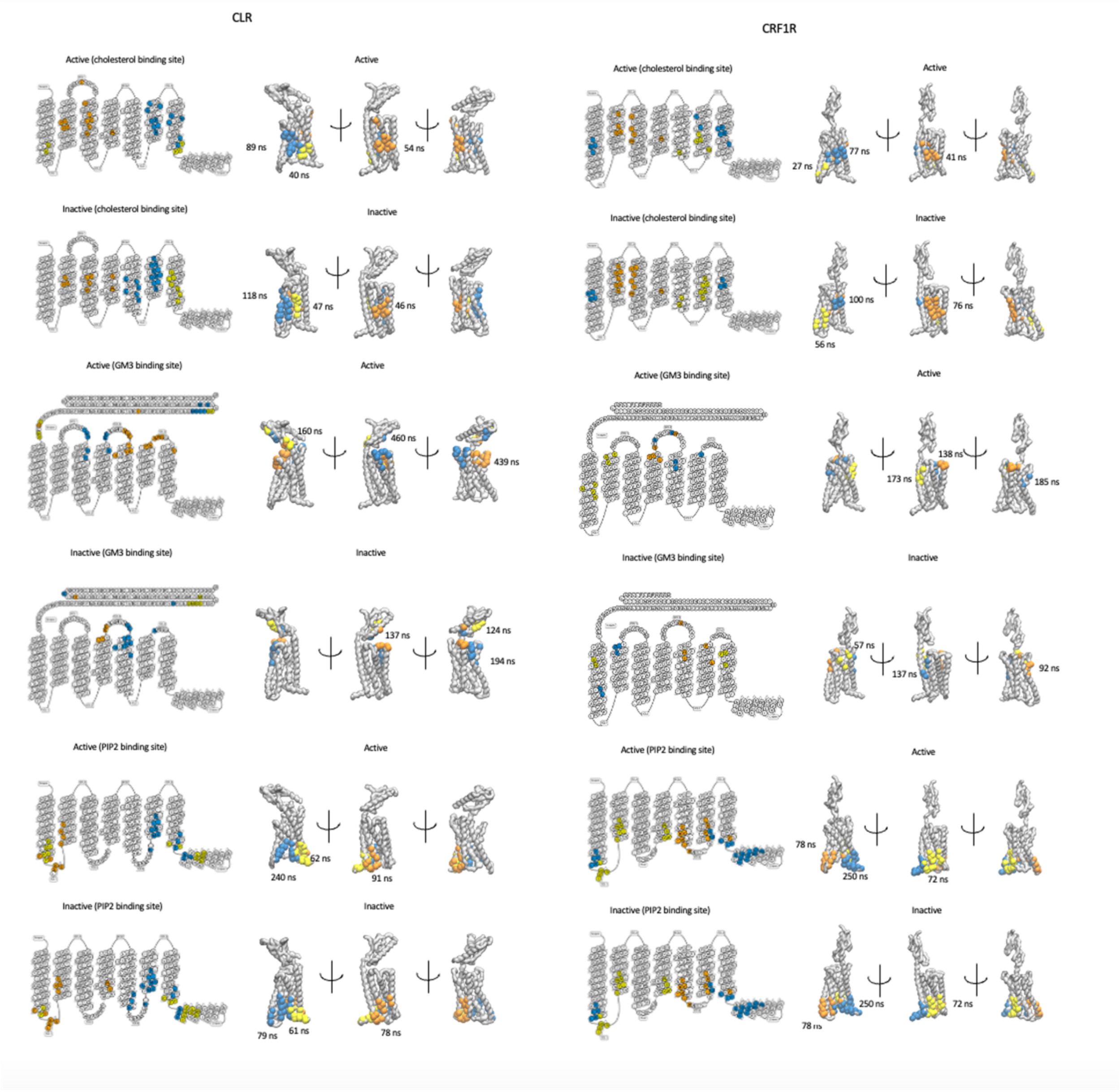

CLR
CRF1R
Active (cholesterol binding site)
Active
89 ns
40 ns
54 ns
27 ns
77 ns
41 ns
Inactive (cholesterol binding site)
Inactive
118 ns
47 ns
46 ns
100 ns
56 ns
76 ns
Active (GM3 binding site)
160 ns
460 ns
439 ns
173 ns
138 ns
185 ns
Inactive (GM3 binding site)
137 ns
124 ns
194 ns
57 ns
137 ns
92 ns
Active (PIP2 binding site)
240 ns
62 ns
91 ns
78 ns
250 ns
72 ns
Inactive (PIP2 binding site)
79 ns
61 ns
78 ns
250 ns
72 ns

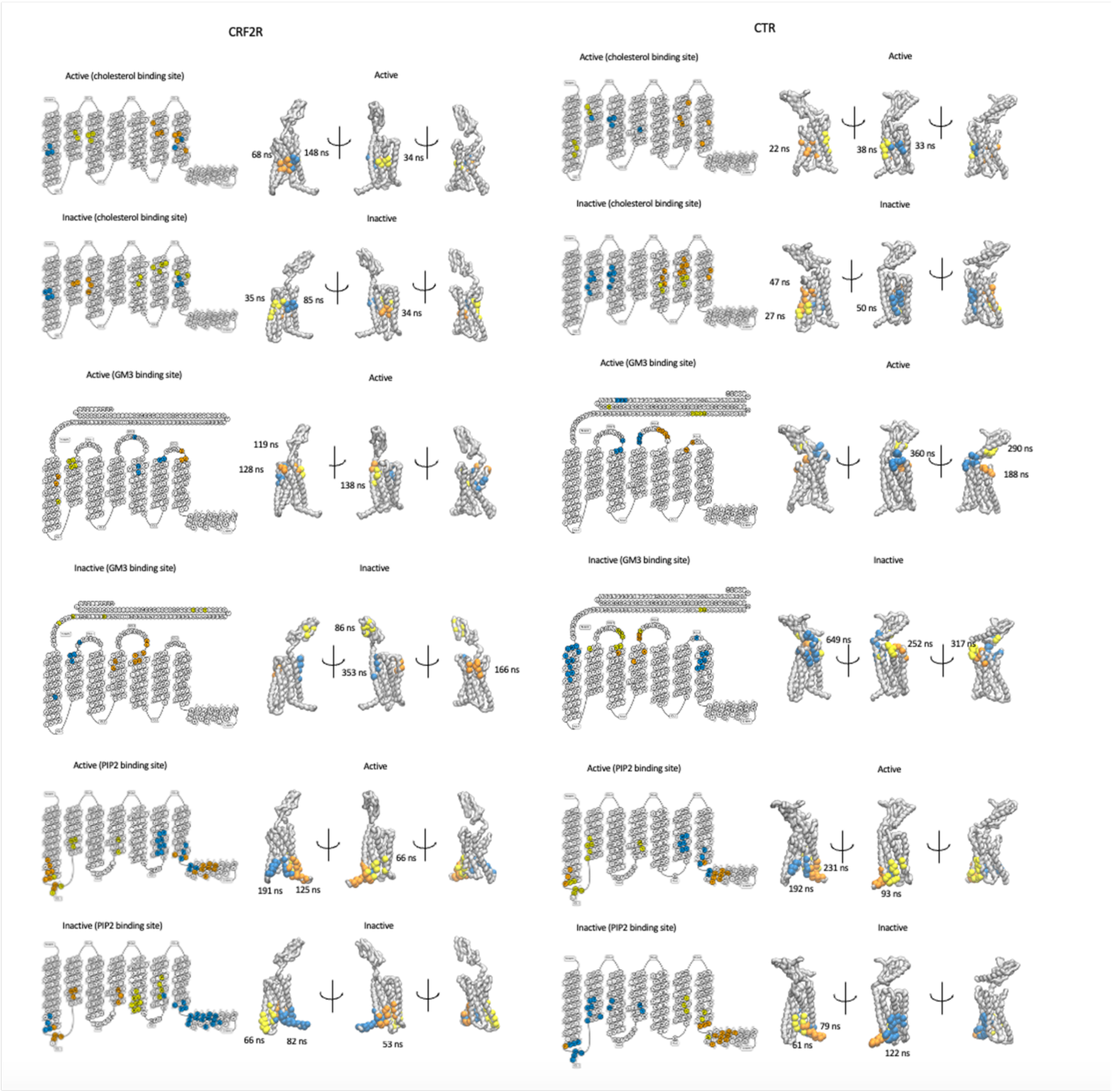

CRF2R
CTR
Active (cholesterol binding site)
Active
68 ns
148 ns
34 ns
22 ns
38 ns
33 ns
Inactive (cholesterol binding site)
Inactive
35 ns
85 ns
34 ns
47 ns
27 ns
50 ns
Active (GM3 binding site)
Active
119 ns
128 ns
138 ns
360 ns
290 ns
188 ns
Inactive (GM3 binding site)
Inactive
86 ns
353 ns
166 ns
649 ns
252 ns
317 ns
Active (PIP2 binding site)
Active
191 ns
125 ns
66 ns
231 ns
192 ns
93 ns
Inactive (PIP2 binding site)
Inactive
66 ns
82 ns
53 ns
79 ns
61 ns
122 ns

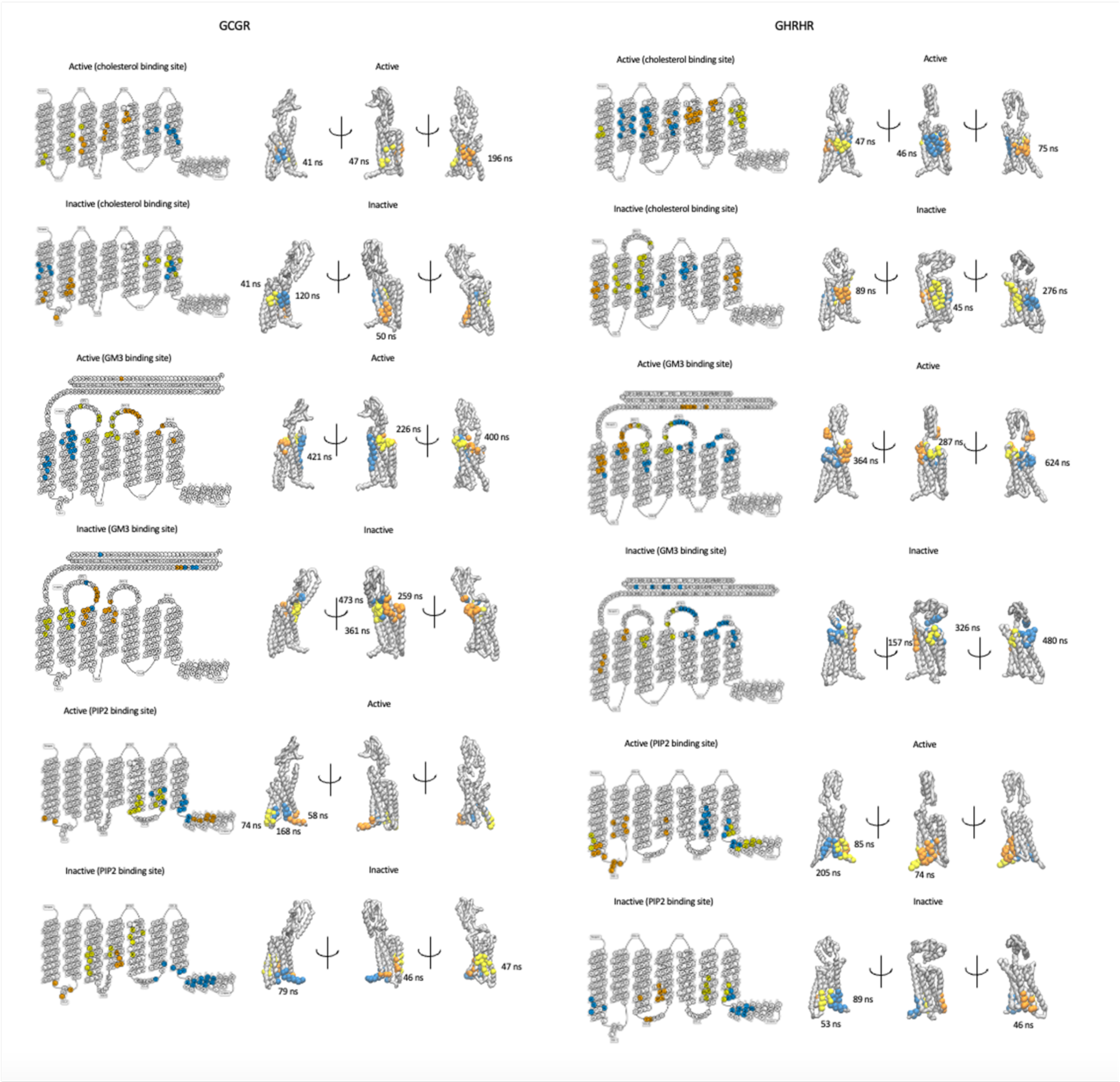

GCGR
GHRHR
Active (cholesterol binding site)
Active
41 ns
47 ns
196 ns
Inactive (cholesterol binding site)
Inactive
41 ns
120 ns
50 ns
Active (GM3 binding site)
Active
421 ns
226 ns
400 ns
Inactive (GM3 binding site)
Inactive
473 ns
361 ns
259 ns
Active (PIP2 binding site)
Active
74 ns
168 ns
58 ns
Inactive (PIP2 binding site)
Inactive
79 ns
46 ns
47 ns
Active (cholesterol binding site)
Active
47 ns
46 ns
75 ns
Inactive (cholesterol binding site)
Inactive
89 ns
45 ns
276 ns
Active (GM3 binding site)
Active
364 ns
287 ns
624 ns
Inactive (GM3 binding site)
Inactive
157 ns
326 ns
480 ns
Active (PIP2 binding site)
Active
85 ns
205 ns
74 ns
Inactive (PIP2 binding site)
Inactive
89 ns
53 ns
46 ns

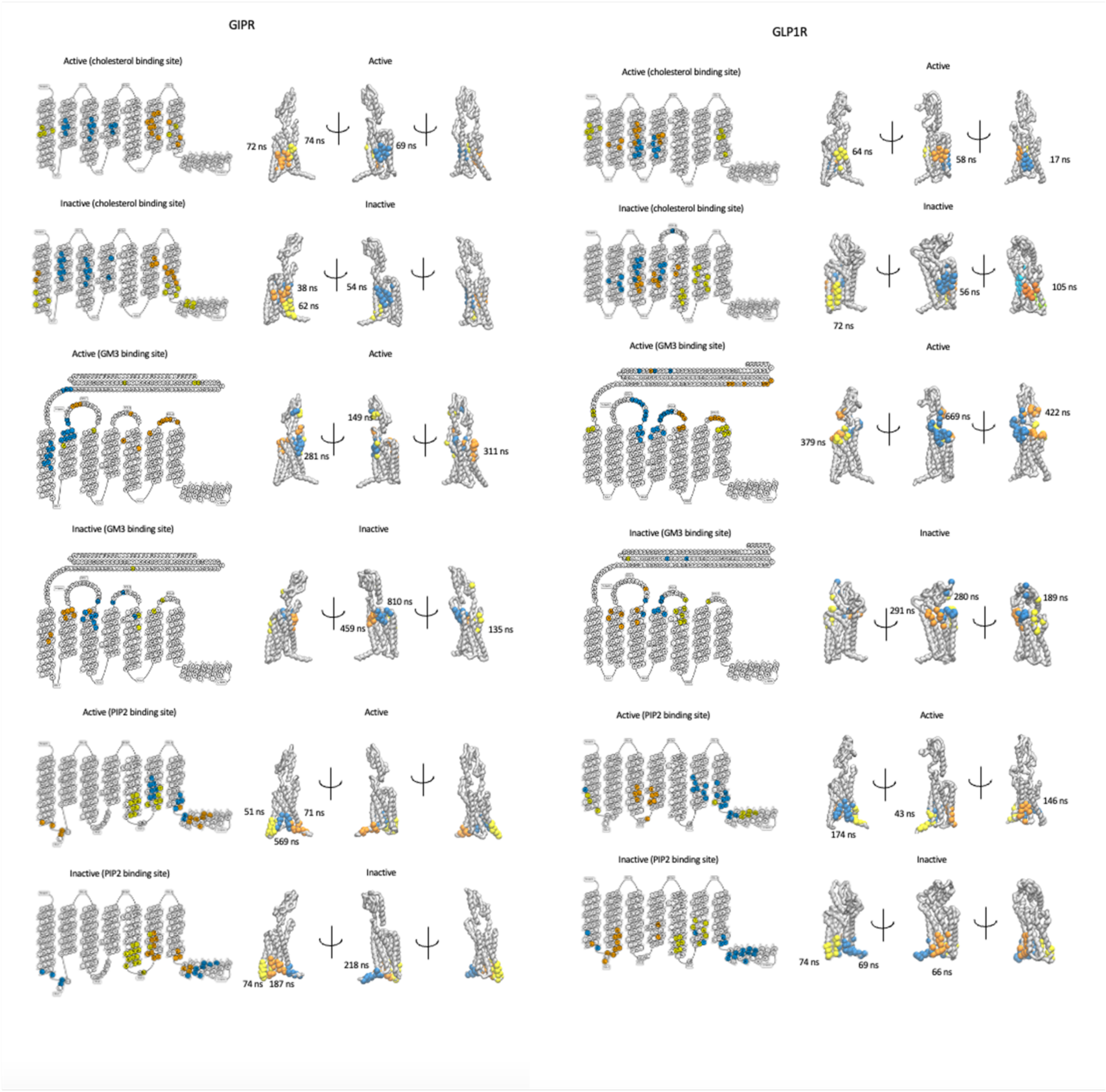
GIPR
Active (cholesterol binding site)
Active
72 ns
74 ns
69 ns
Inactive (cholesterol binding site)
Inactive
38 ns
62 ns
54 ns
Active (GM3 binding site)
Active
149 ns
281 ns
311 ns
Inactive (GM3 binding site)
Inactive
810 ns
459 ns
135 ns
Active (PIP2 binding site)
Active
51 ns
71 ns
569 ns
Inactive (PIP2 binding site)
Inactive
74 ns
187 ns
218 ns
GLP1R
Active (cholesterol binding site)
Active
64 ns
58 ns
17 ns
Inactive (cholesterol binding site)
Inactive
56 ns
105 ns
72 ns
Active (GM3 binding site)
Active
669 ns
422 ns
379 ns
Inactive (GM3 binding site)
Inactive
291 ns
280 ns
189 ns
Active (PIP2 binding site)
Active
146 ns
43 ns
174 ns
Inactive (PIP2 binding site)
Inactive
74 ns
69 ns
66 ns

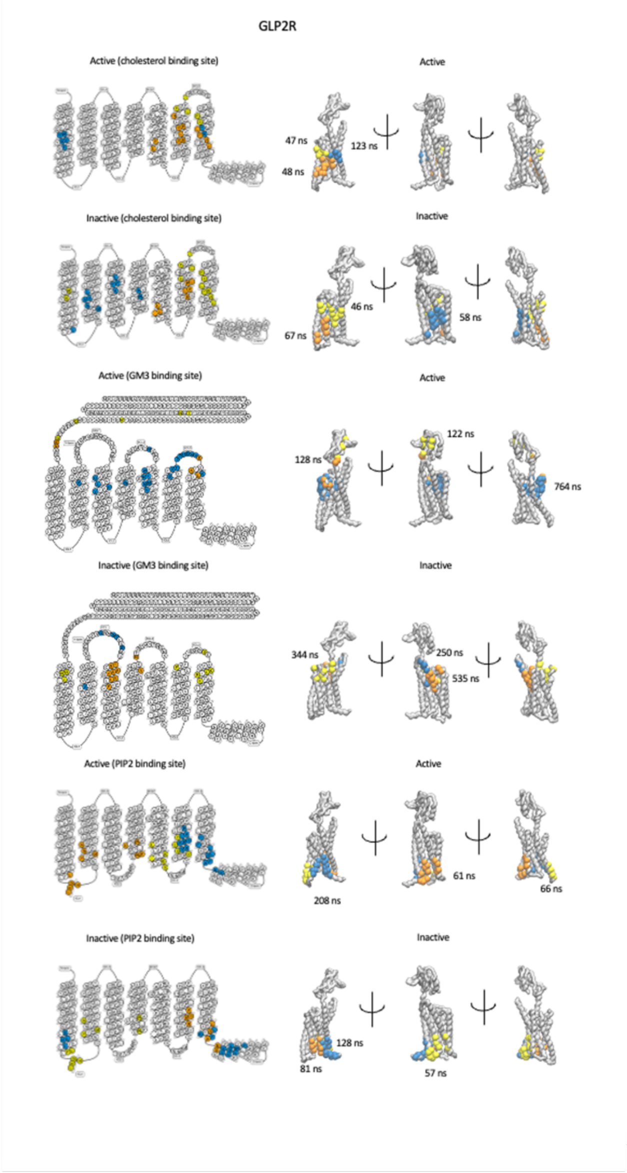
GLP2R
Active (cholesterol binding site)
Active
47 ns
48 ns
123 ns
Inactive (cholesterol binding site)
Inactive
67 ns
46 ns
58 ns
Active (GM3 binding site)
Active
128 ns
122 ns
764 ns
Inactive (GM3 binding site)
Inactive
344 ns
250 ns
535 ns
Active (PIP2 binding site)
Active
208 ns
61 ns
66 ns
Inactive (PIP2 binding site)
Inactive
81 ns
128 ns
57 ns

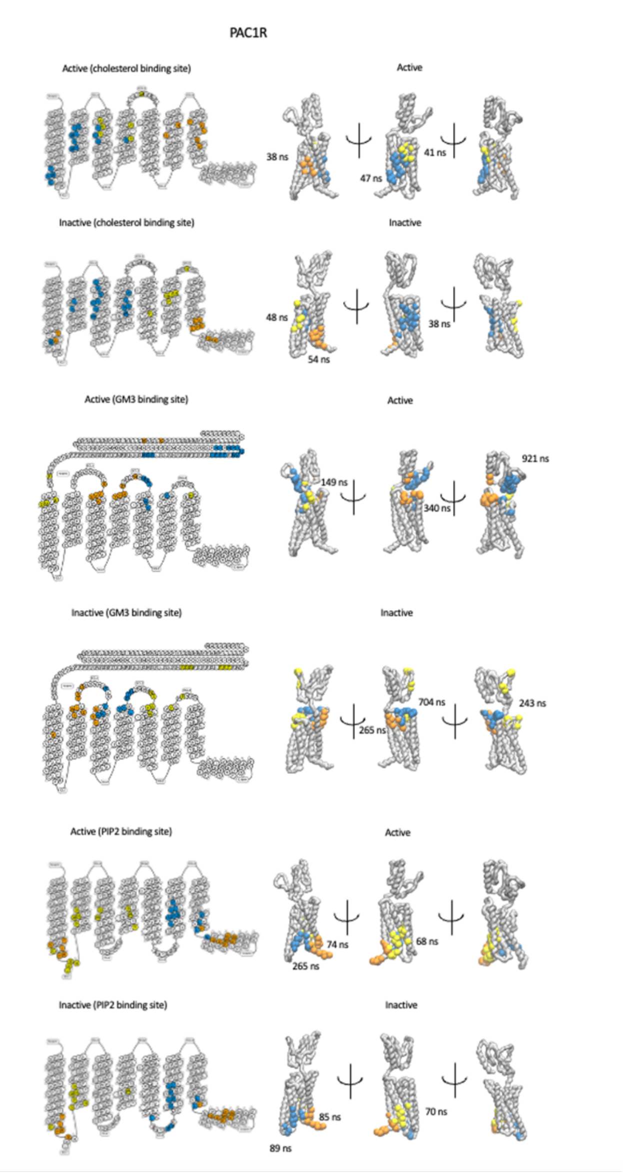
PAC1R
Active (cholesterol binding site)
Active
38 ns
47 ns
41 ns
Inactive (cholesterol binding site)
Inactive
48 ns
54 ns
38 ns
Active (GM3 binding site)
Active
149 ns
340 ns
921 ns
Inactive (GM3 binding site)
Inactive
265 ns
704 ns
243 ns
Active (PIP2 binding site)
Active
265 ns
74 ns
68 ns
Inactive (PIP2 binding site)
Inactive
89 ns
85 ns
70 ns

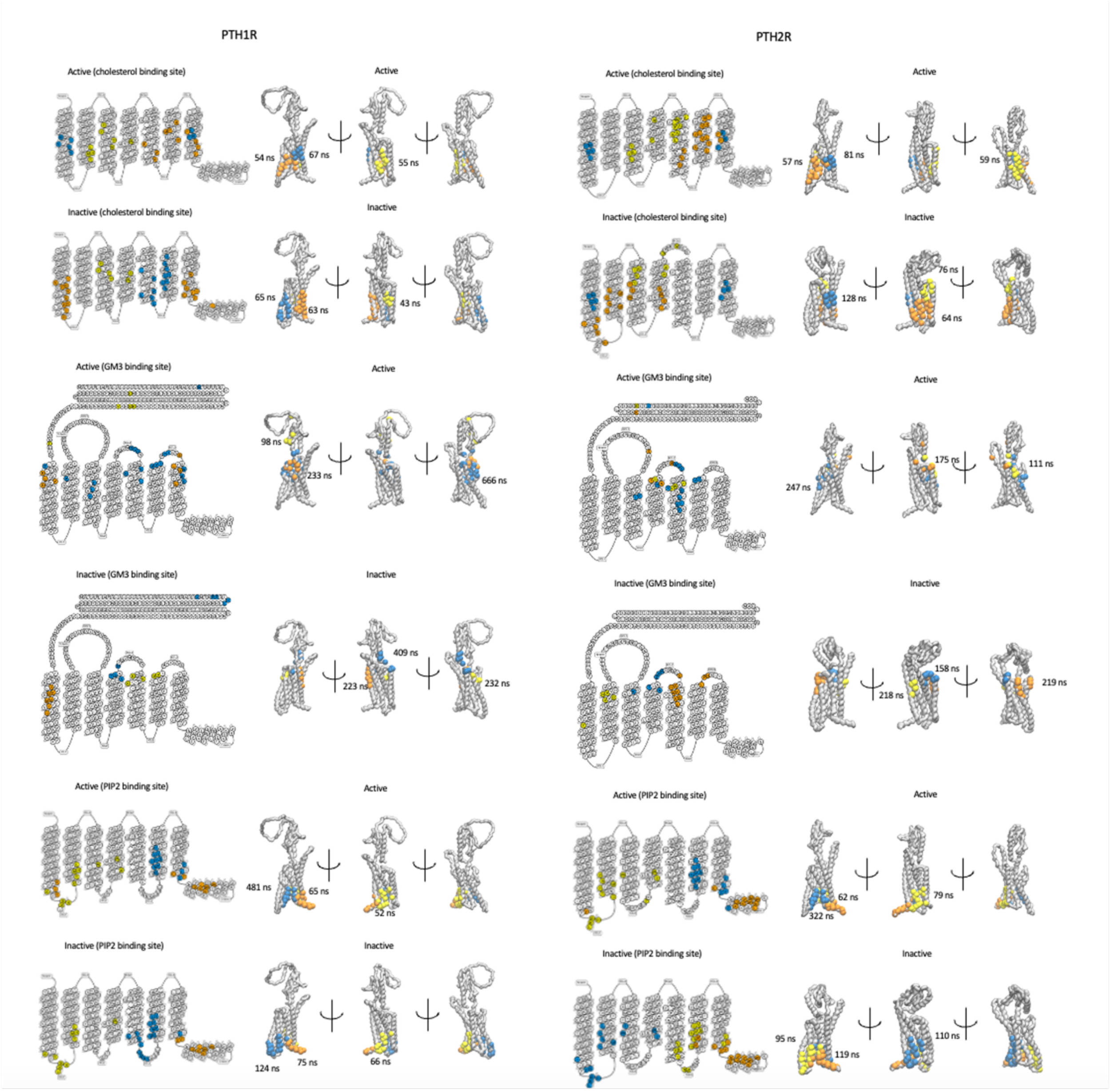
PTH1R
Active (cholesterol binding site)
Active
54 ns
67 ns
55 ns
Inactive (cholesterol binding site)
Inactive
65 ns
63 ns
43 ns
Active (GM3 binding site)
Active
98 ns
233 ns
666 ns
Inactive (GM3 binding site)
Inactive
409 ns
223 ns
232 ns
Active (PIP2 binding site)
Active
481 ns
65 ns
52 ns
Inactive (PIP2 binding site)
Inactive
124 ns
75 ns
66 ns
PTH2R
Active (cholesterol binding site)
Active
57 ns
81 ns
59 ns
Inactive (cholesterol binding site)
Inactive
128 ns
76 ns
64 ns
Active (GM3 binding site)
Active
247 ns
175 ns
111 ns
Inactive (GM3 binding site)
Inactive
218 ns
158 ns
219 ns
Active (PIP2 binding site)
Active
62 ns
322 ns
79 ns
Inactive (PIP2 binding site)
Inactive
95 ns
119 ns
110 ns

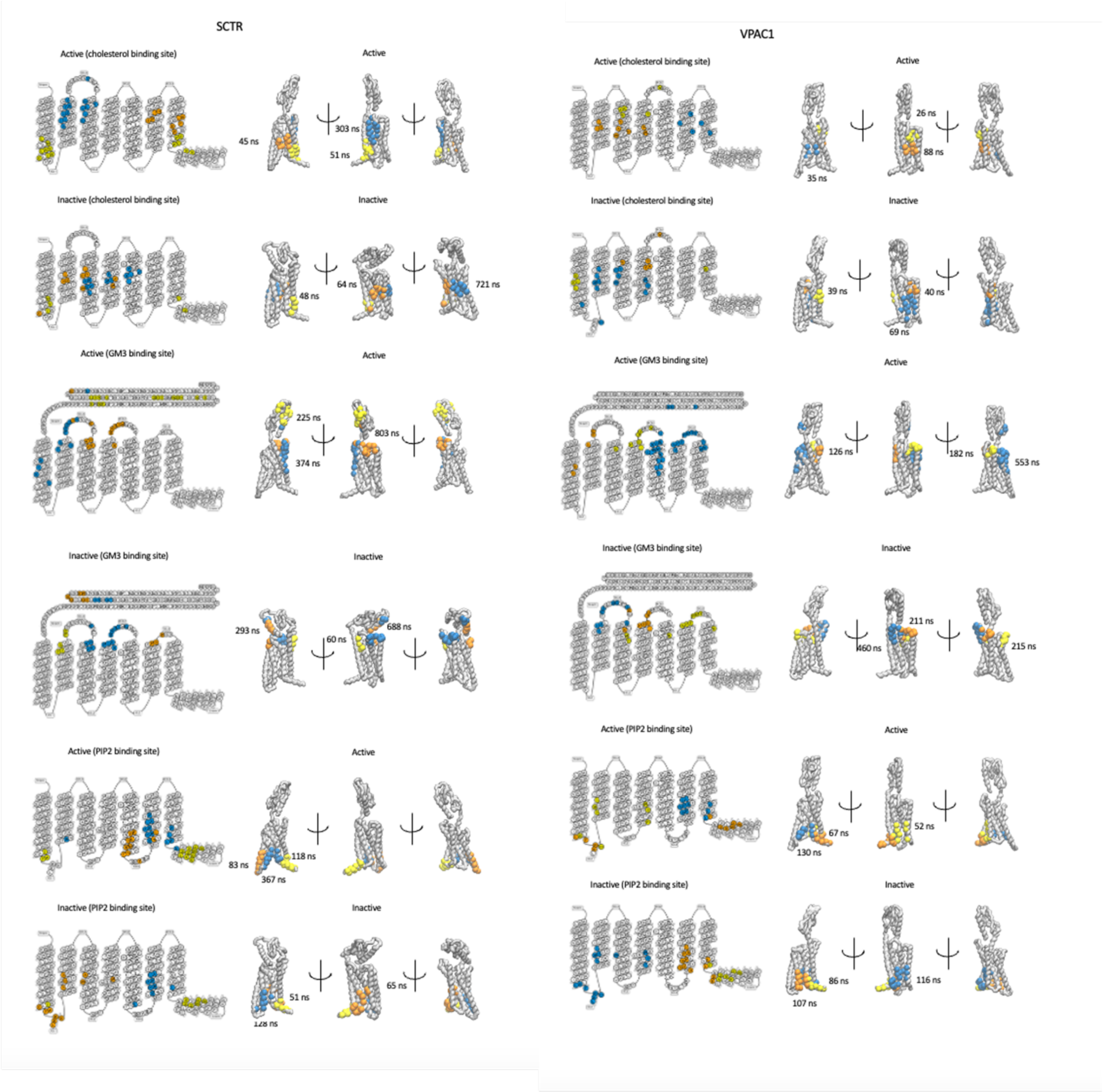
SCTR
Active (cholesterol binding site)
Active
45 ns
303 ns
51 ns
Inactive (cholesterol binding site)
Inactive
48 ns
64 ns
721 ns
Active (GM3 binding site)
Active
225 ns
374 ns
803 ns
Inactive (GM3 binding site)
Inactive
293 ns
60 ns
688 ns
Active (PIP2 binding site)
Active
83 ns
118 ns
367 ns
Inactive (PIP2 binding site)
Inactive
51 ns
128 ns
65 ns
VPAC1
Active (cholesterol binding site)
Active
35 ns
26 ns
88 ns
Inactive (cholesterol binding site)
Inactive
39 ns
40 ns
69 ns
Active (GM3 binding site)
Active
126 ns
182 ns
553 ns
Inactive (GM3 binding site)
Inactive
460 ns
211 ns
215 ns
Active (PIP2 binding site)
Active
67 ns
130 ns
52 ns
Inactive (PIP2 binding site)
Inactive
86 ns
107 ns
116 ns

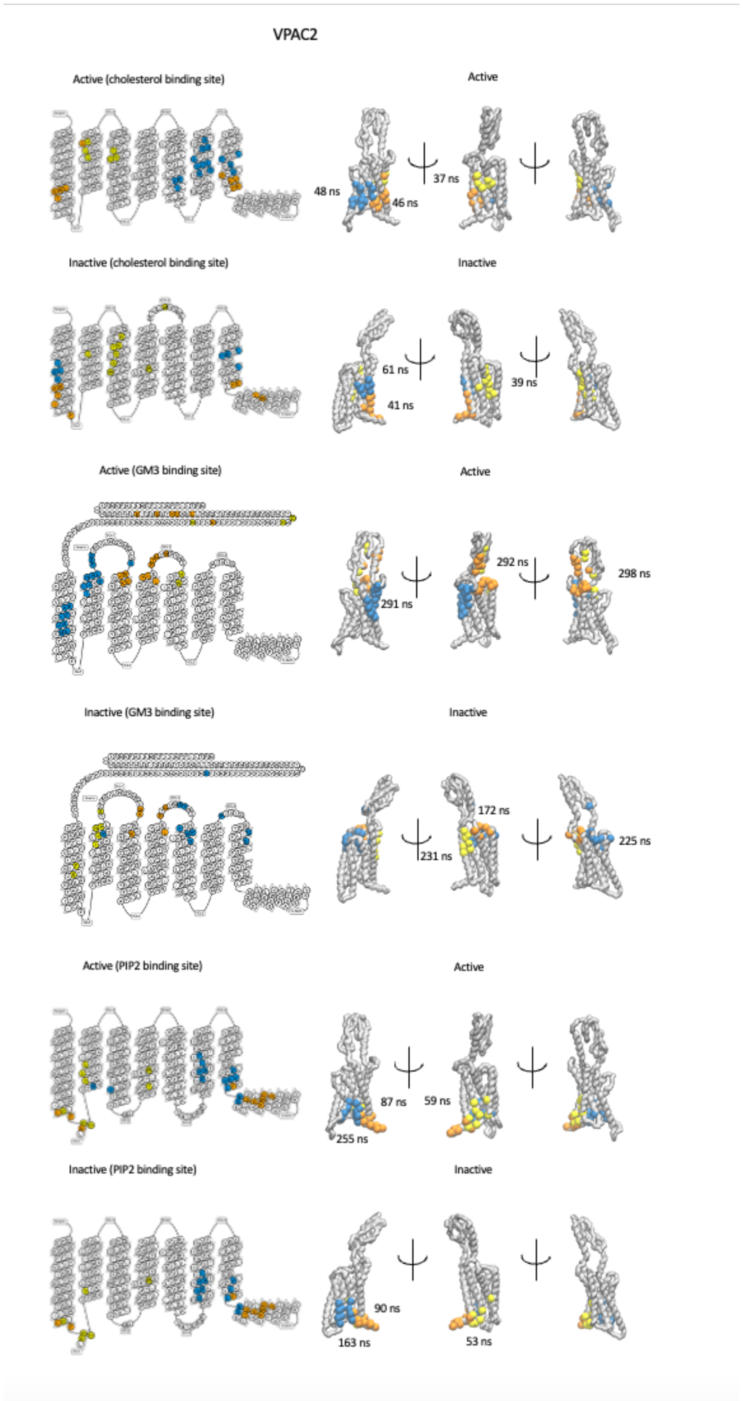


**Fig. S10.**

**Cholesterol, $PIP_2$ and GM3 binding sites for class B1 receptors**. The top 30 highest occupancy values for cholesterol, GM3 and $PIP_2$ are mapped to the GPCR snakeplot and 3D structure for active and inactive states. The top three binding sites calculated in terms of residence time are shown on the snakeplot and 3D structure. Site I (cyan), Site II (orange), and Site III (yellow/lime). The averaged residence time for each site is shown.

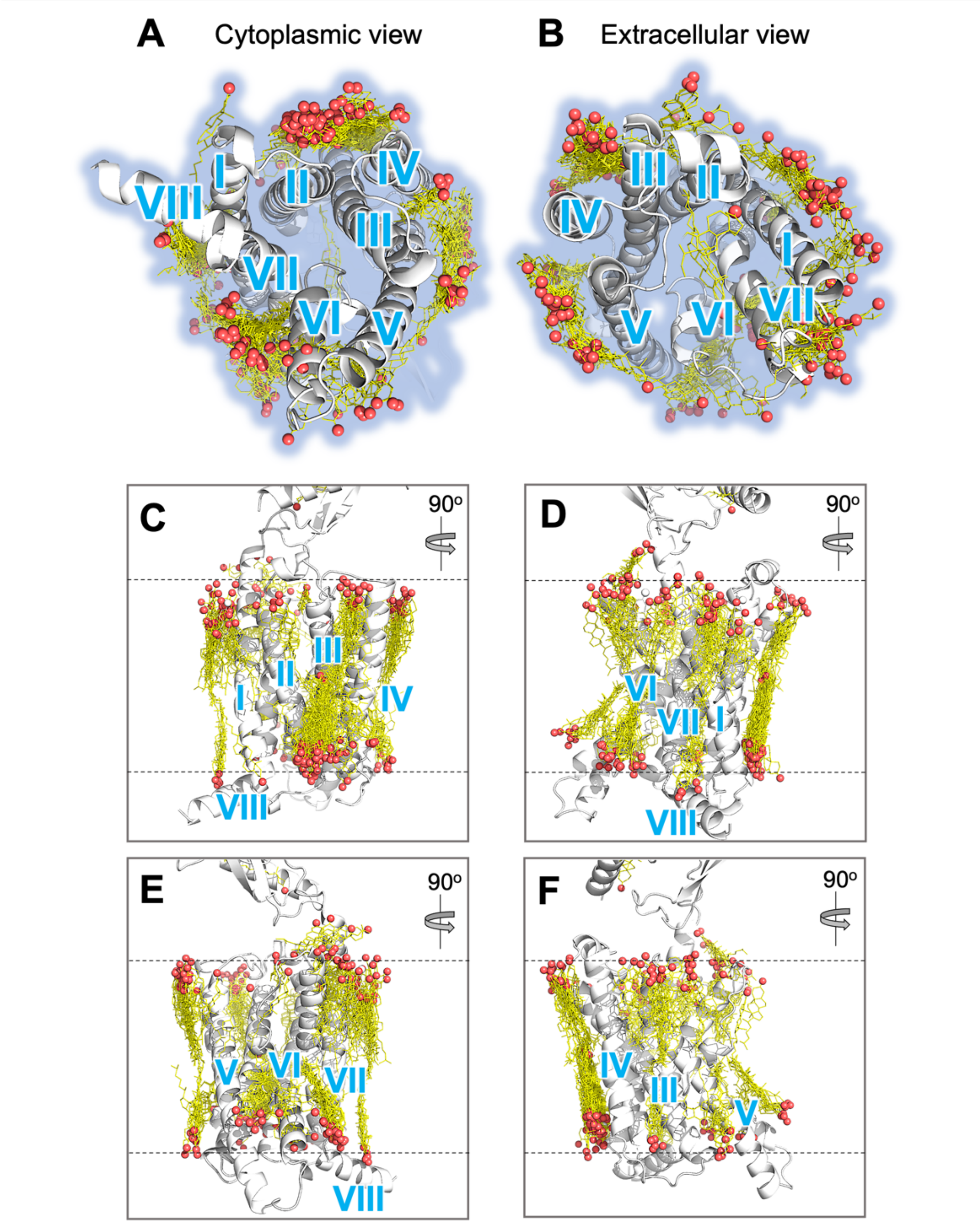


**Fig. S11.**

**Overlay of 225 AI predicted cholesterol interaction sites across all 15 Class B1 GPCRs.** Cholesterol molecules are shown as yellow sticks, with the position of the oxygen atom in hydroxyl headgroup denoted by a red sphere. Transmembrane helices are labelled in blue. For each receptor five Chai-1 AI models were generated, with each model containing three cholesterols poses. Models were created using the same GPCR primary sequence used in MD simulations, and the SMILES string for cholesterol into the Chai-1 web interface (https://lab.chaidiscovery.com/), with the multiple sequence alignment option active. See Methods for further details of AI model generation.

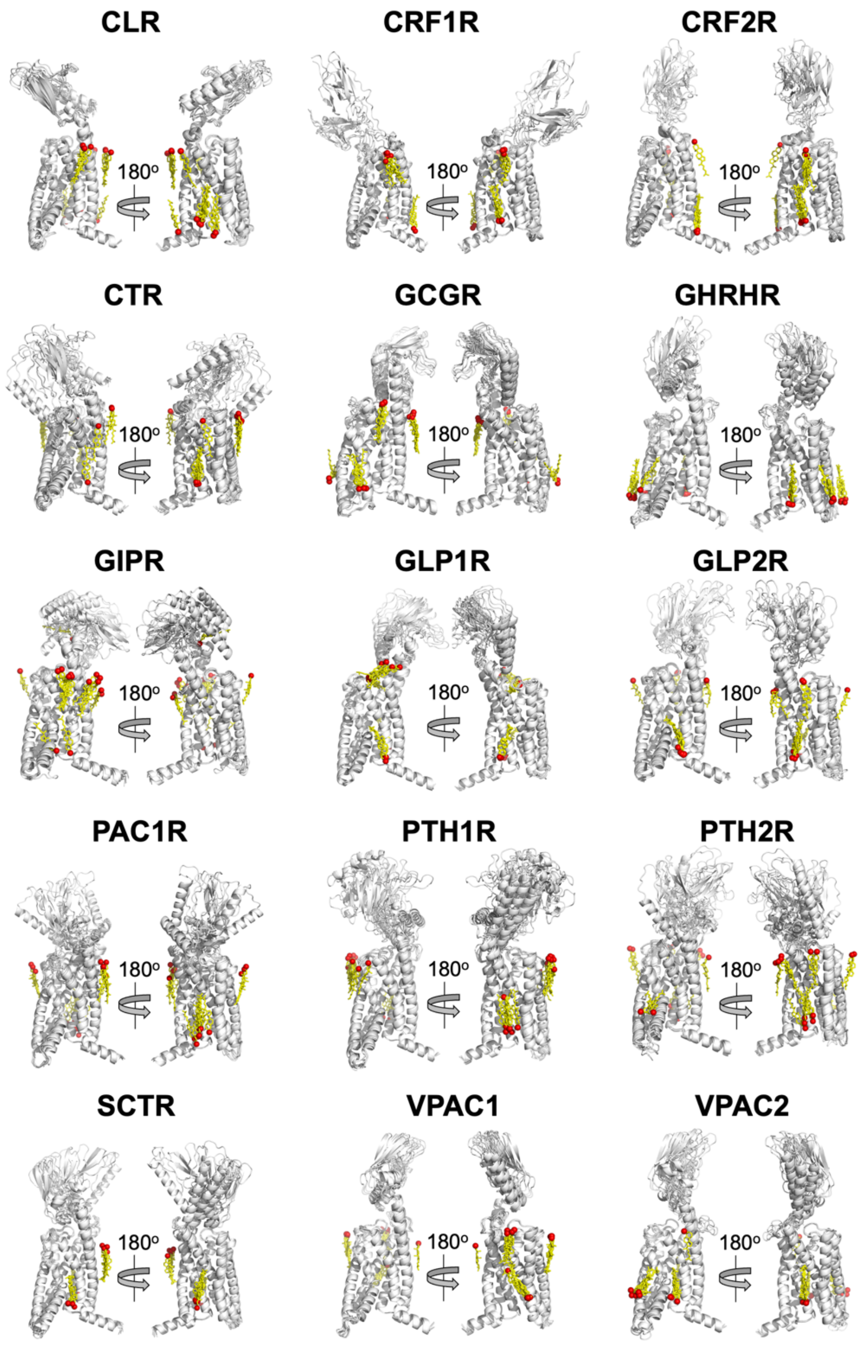


**Fig. S12.**

**AI predicted cholesterol interaction sites on all 15 Class B1 GPCRs**. Cholesterol molecules are shown as yellow sticks, with the position of the oxygen atom in hydroxyl headgroup denoted by a red sphere. For each receptor five AI models were generated, with each model containing three cholesterols poses. These output models are shown overlaid for each Class B1 receptor. See Methods for further details of Chai-1 model generation.

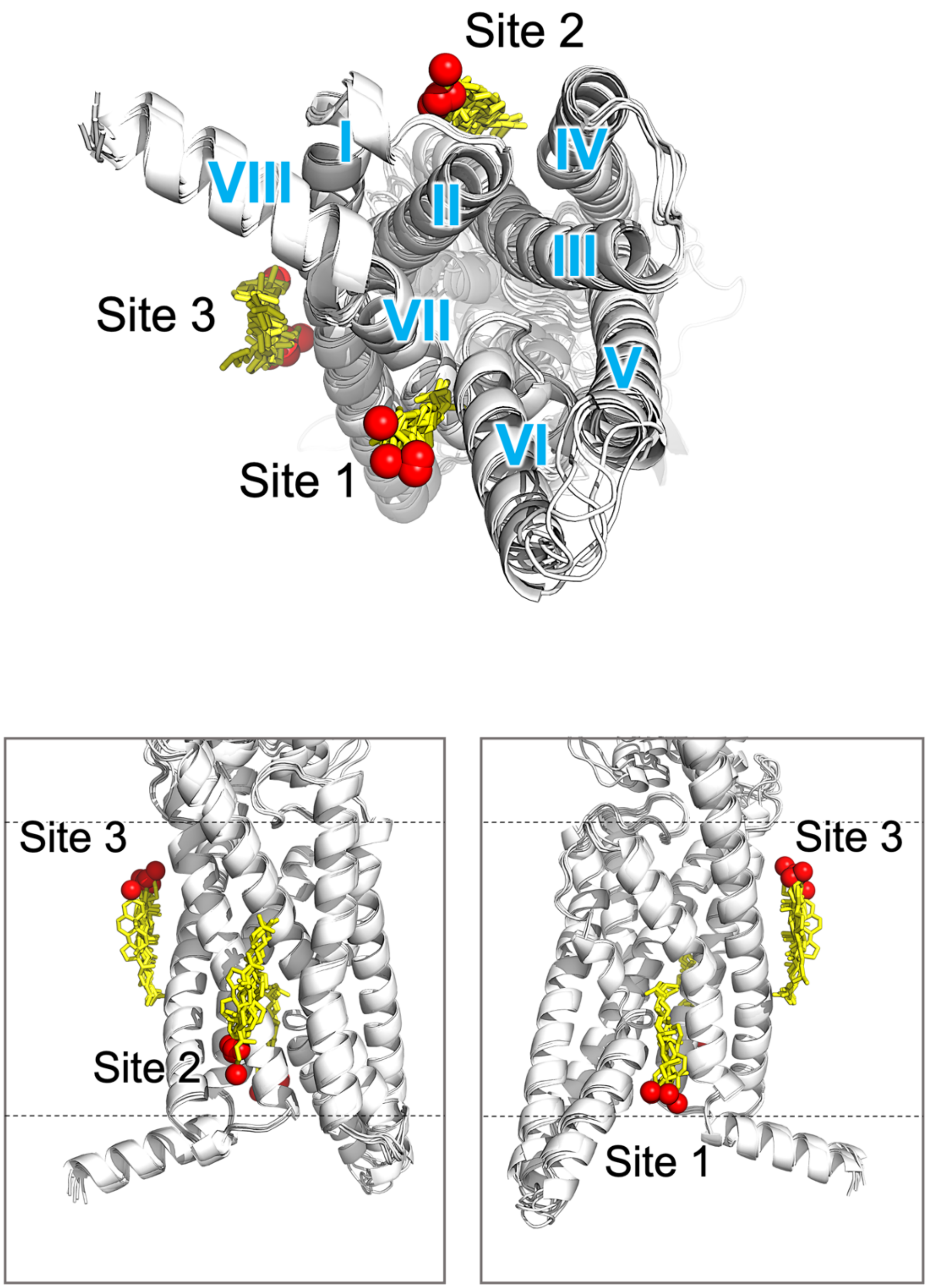


**Fig. S13.**

**AI predicted cholesterol interaction sites on the Sec receptor (SCTR).** Cholesterol molecules are shown as yellow sticks, with the position of the oxygen atom in hydroxyl headgroup denoted by a red sphere. Five Chai-1 models are shown overlaid, each containing three cholesterol molecules. See Methods for further details of Chai-1 model generation.

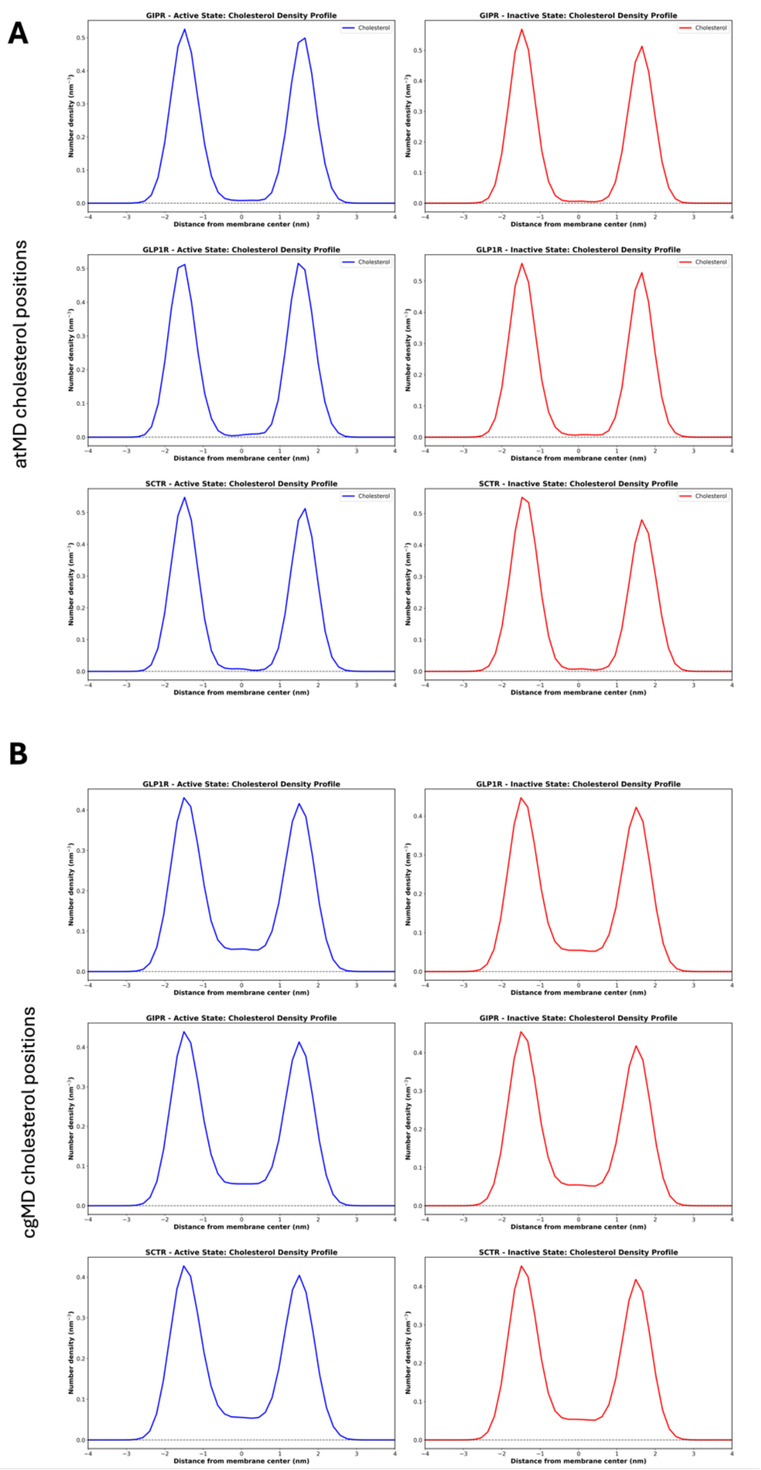


**Fig. S14.**

**Comparison of cholesterol headgroup position in cgMD vs atomistic simulations.** A) Calculated cholesterol headgroup bead Z position (membrane normal) atMD simulations of active and inactive states (averaged across three repeats) B) As for (A) but using the cg ROH particle for cholesterol.

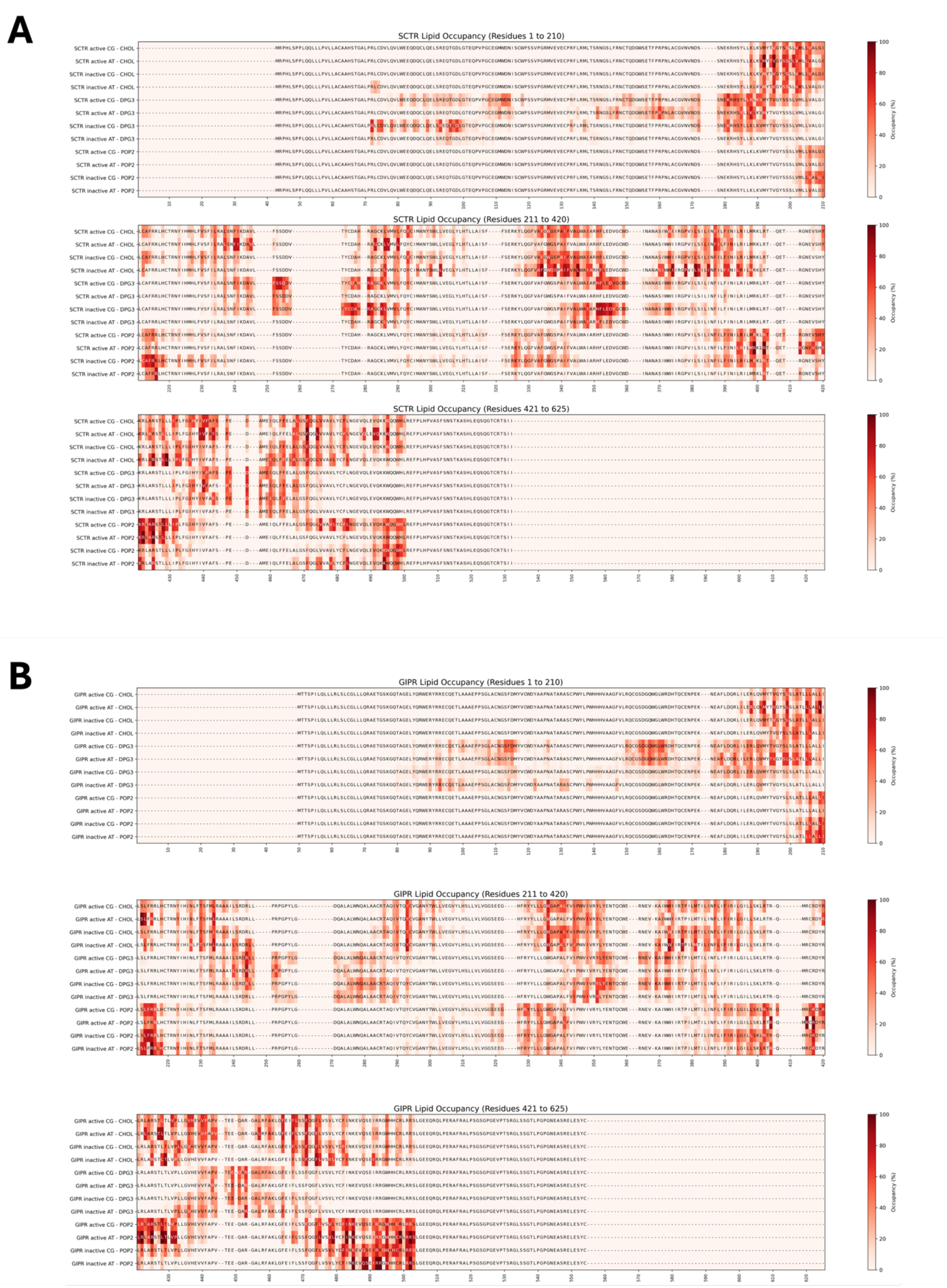
A
SCTR Lipid Occupancy (Residues 1 to 210)
SCTR Lipid Occupancy (Residues 211 to 420)
SCTR Lipid Occupancy (Residues 421 to 625)
SCTR active CG - CHOL
SCTR active AT - CHOL
SCTR inactive CG - CHOL
SCTR inactive AT - CHOL
SCTR active CG - DPG3
SCTR active AT - DPG3
SCTR inactive CG - DPG3
SCTR inactive AT - DPG3
SCTR active CG - POP2
SCTR active AT - POP2
SCTR inactive CG - POP2
SCTR inactive AT - POP2
Occupancy (%)
B
GIPR Lipid Occupancy (Residues 1 to 210)
GIPR Lipid Occupancy (Residues 211 to 420)
GIPR Lipid Occupancy (Residues 421 to 625)
GIPR active CG - CHOL
GIPR active AT - CHOL
GIPR inactive CG - CHOL
GIPR inactive AT - CHOL
GIPR active CG - DPG3
GIPR active AT - DPG3
GIPR inactive CG - DPG3
GIPR inactive AT - DPG3
GIPR active CG - POP2
GIPR active AT - POP2
GIPR inactive CG - POP2
GIPR inactive AT - POP2
Occupancy (%)

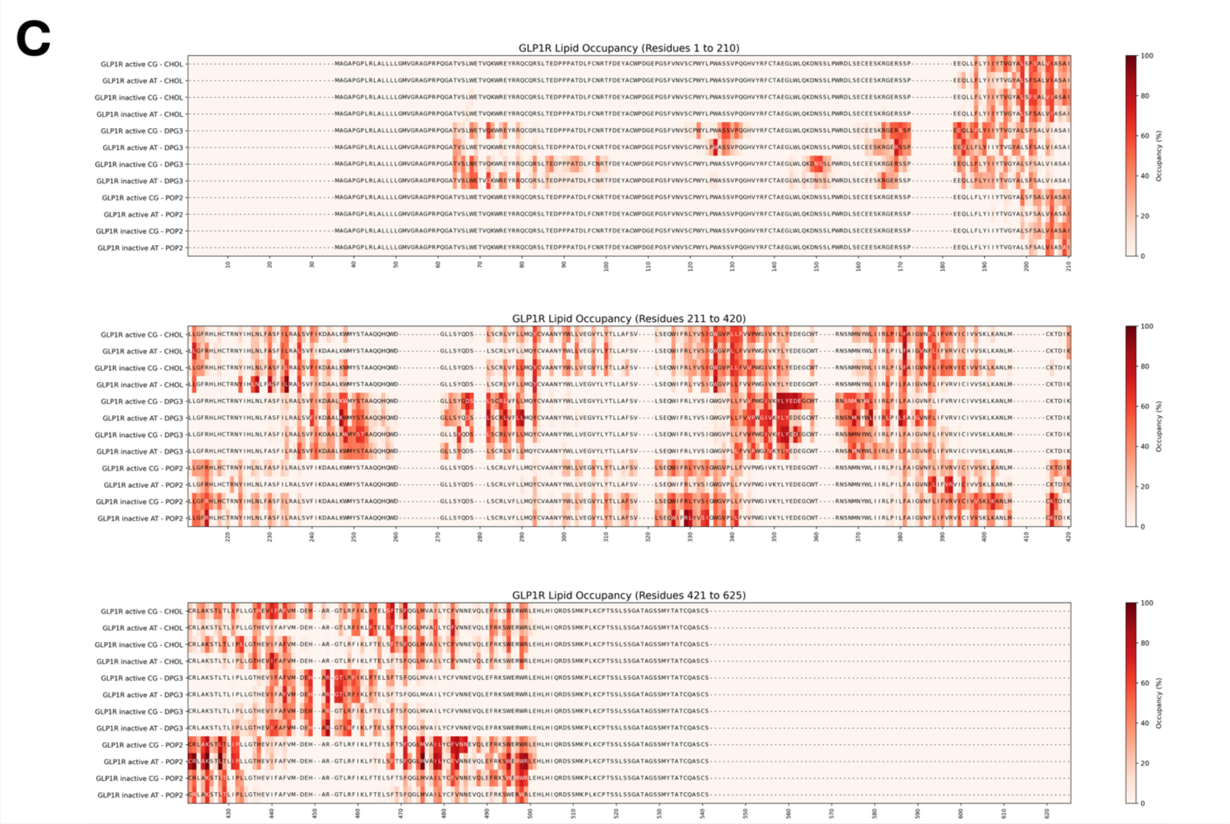


**Fig. S15.**

**Cholesterol, GM2 and PIP₂ interactions in cgMD vs atomistic simulations.** Lipid occupancy (CHOL, GM3 and $PIP_2$) shown as sequence-based heatmaps for cgMD and atomistic simulations of (A) SCTR, (B) GIPR, and (C) GLP1R in both active and inactive states (high occupancy to low occupancy: red to white). The CG and AT data are shown side-by-side for each condition to allow direct comparison (cutoff distances: CG = 7 Å, AT = 4 Å). Each heatmap represents averaged data from three simulation repeats. Martini3 naming: POP2 = $PIP_2$, DPG3 = GM3.

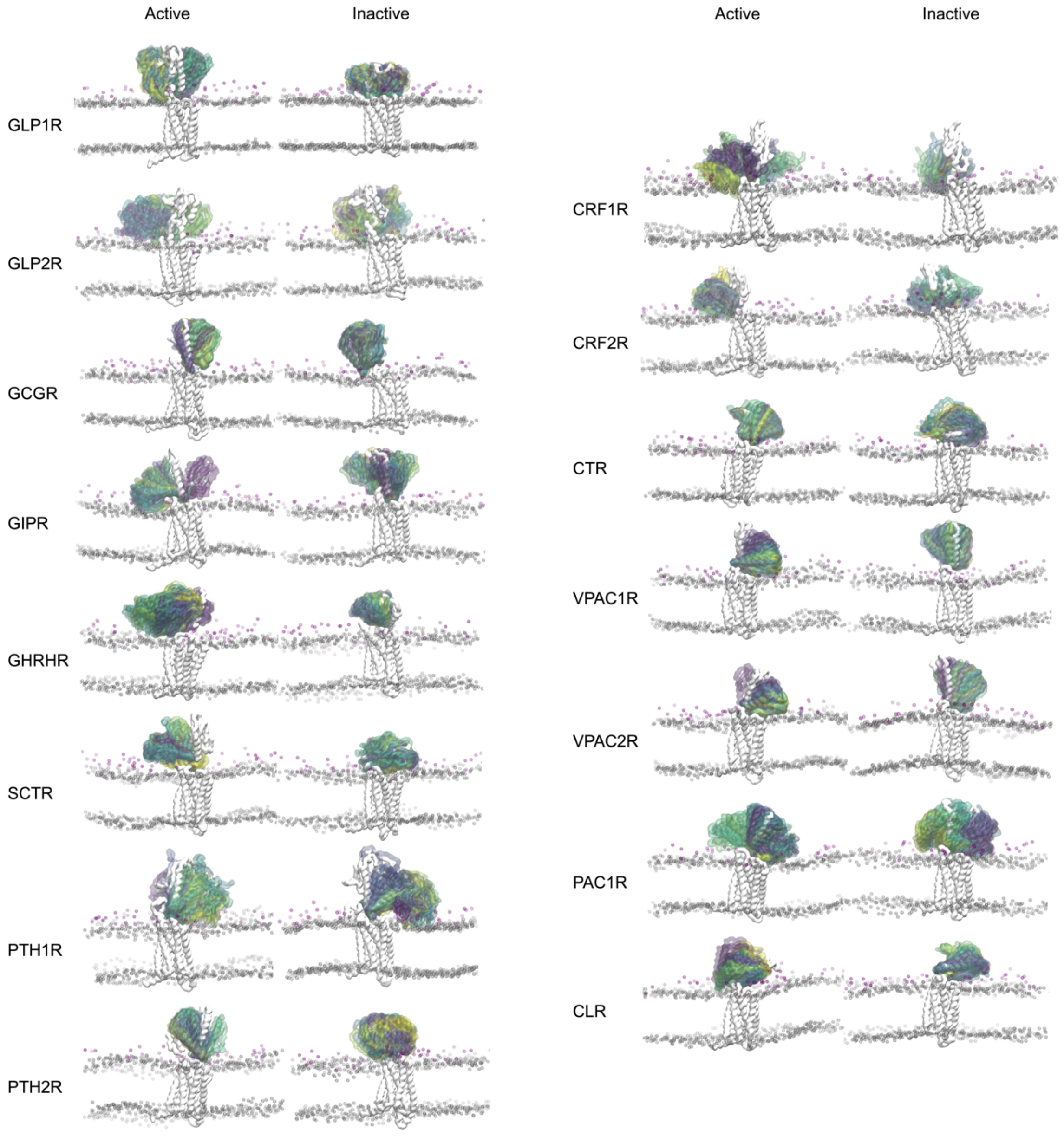


**Fig. S16.**

**ECD motions of class B1 receptors in model plasma membranes.** The starting structure is shown as solid grey, whilst the ECD motion relative to the TM bundle is shown coloured by timestep (0 to 10 us, blue-green-yellow). Phospholipid headgroup beads are shown as transparent grey spheres and GM3 GL beads are shown in magenta. Data is shown for the first repeat for each receptor state.

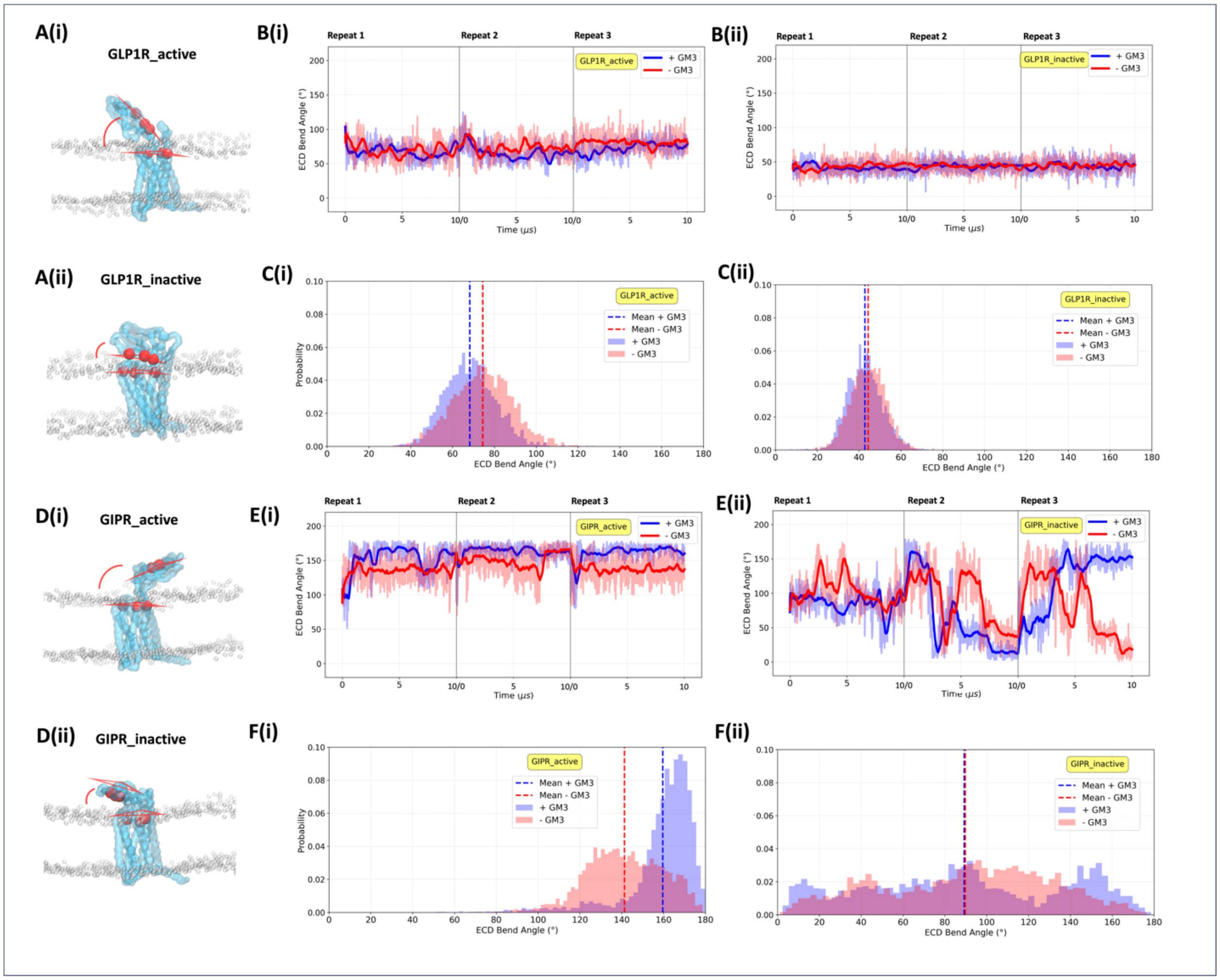


**Fig. S17.**

Effect of GM3 on ECD motion of GLP1R and GIPR (**A**) (i) and (ii) show the angle between the two planes formed by three residues (Y42, C46, T65, and D198, R227, K288) in GLP-1R, measured throughout the cgMD simulations. (**B**) The bend angle is plotted with a rolling average for the three repeats in the (i) active state and (ii) inactive state (**C**) Bend angle histograms for the (i) active state and (ii) inactive state. (**D**) (i) and (ii) show the bend angle between the two planes formed by three residues (Y42, C46, C70, and D191, R217, R278) in GIPR. (**E**) Bend angle over time for active (i) and inactive states (ii). (**F**) Histograms of bend angles. Simulations with 10% GM3 are shown in blue, simulations with 0% GM3 are shown in red throughout.

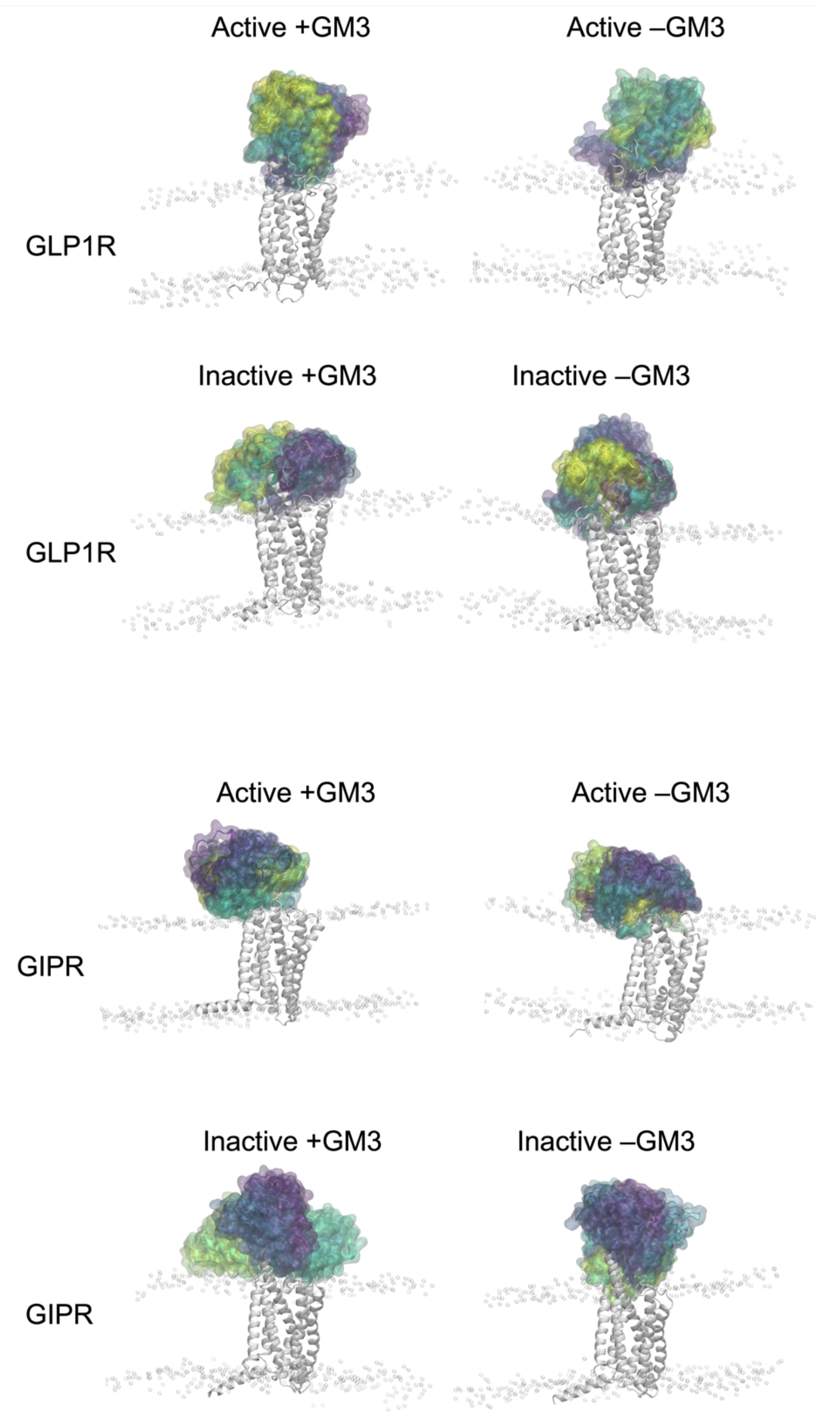


**Fig. S18.**

**ECD motions of GLP1R and GIPR at atomistic resolution.** The ECD motion relative to the TM bundle is shown coloured by timestep (replicate 1, replicate 2, replicate 3, blue-green-yellow). Phospholipid headgroup beads are shown as transparent grey spheres. Left hand panels: membranes included GM3, right hand panels: GM3 absent.

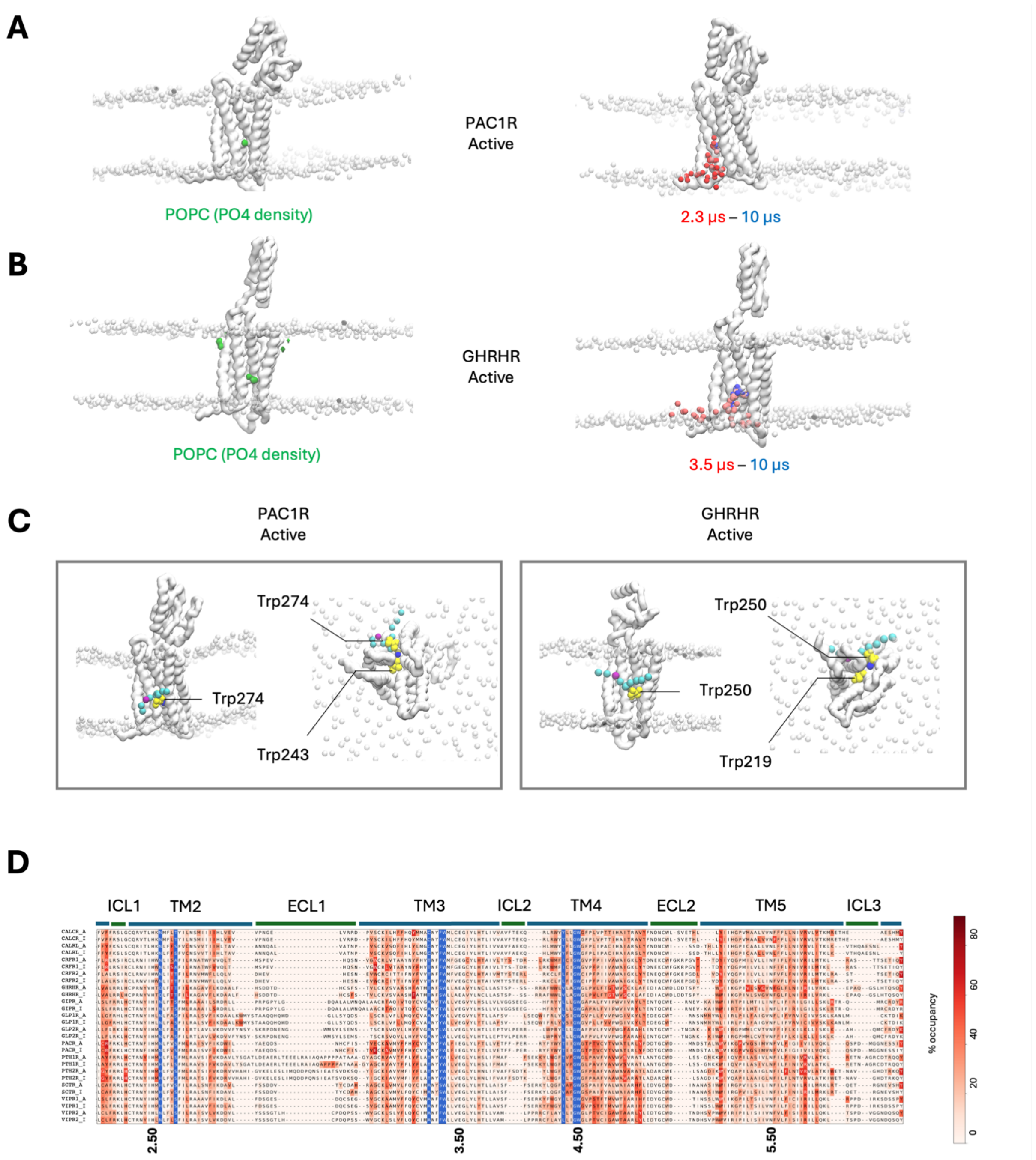


**Fig. S19.**

**Rare deep membrane binding events of POPC in PAC1R and GHRHR active states. A)** Volumetric density of POPC PO4 headgroups across 3 x 10 μs repeats. Timestep series showing headgroup positions of deep binding POPC molecule during one replicate. **B)** As in A) but GHRHR. **C)** Key interactions are mediated by surface and buried Trp residues for both receptors. **D)** Sequence alignment across class B1 GPCRs, with the residues involved in the POPC binding site highlighted in blue.

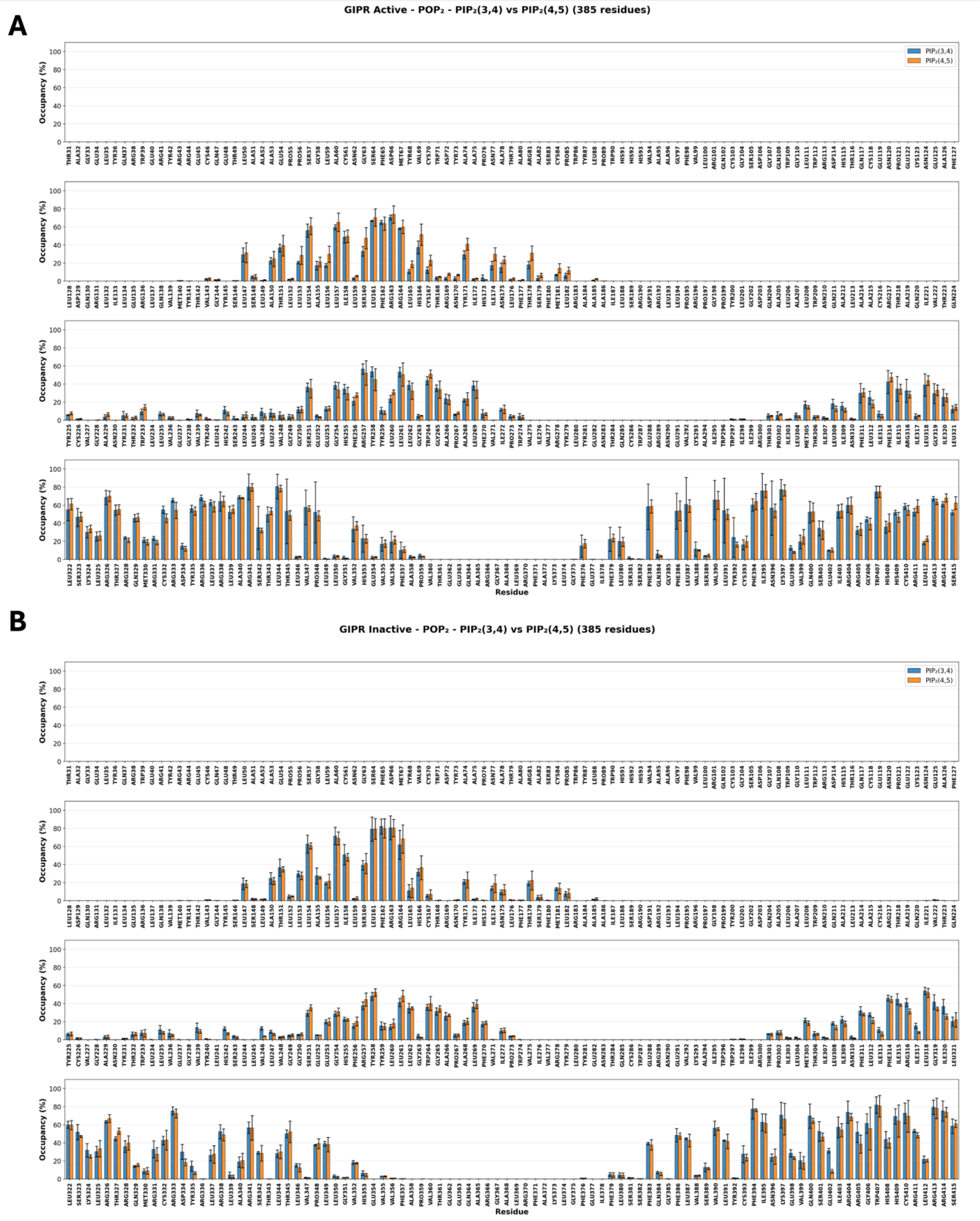
A
GIPR Active - POP₂ - PIP₂(3,4) vs PIP₂(4,5) (385 residues)
PIP₂(3,4)
PIP₂(4,5)
Occupancy (%)
Residue
B
GIPR Inactive - POP₂ - PIP₂(3,4) vs PIP₂(4,5) (385 residues)
PIP₂(3,4)
PIP₂(4,5)
Occupancy (%)
Residue

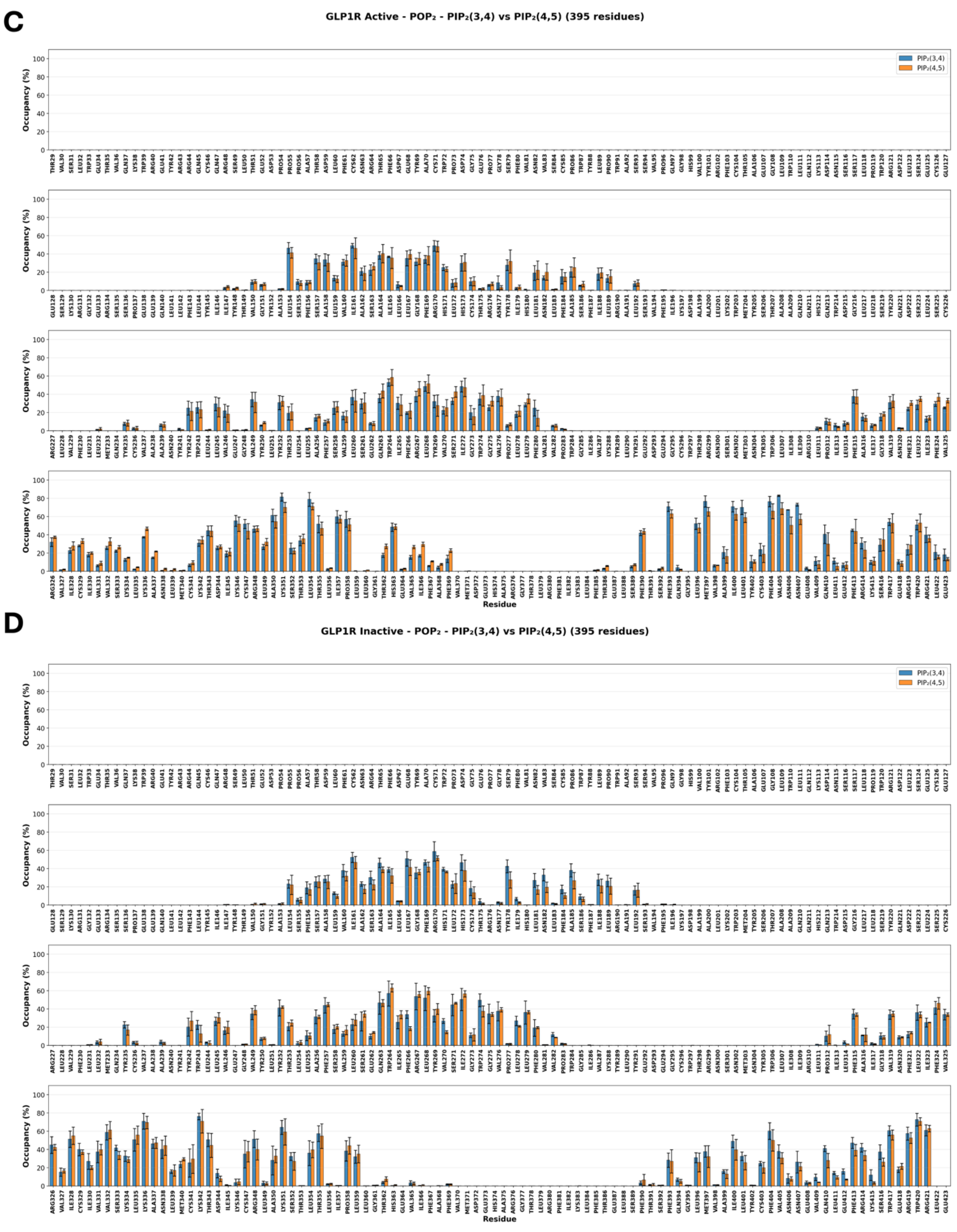
C
GLP1R Active - POP$_2$ - PIP$_2$(3,4) vs PIP$_2$(4,5) (395 residues)
PIP$_2$(3,4)
PIP$_2$(4,5)
Occupancy (%)
Residue
D
GLP1R Inactive - POP$_2$ - PIP$_2$(3,4) vs PIP$_2$(4,5) (395 residues)
PIP$_2$(3,4)
PIP$_2$(4,5)
Occupancy (%)
Residue

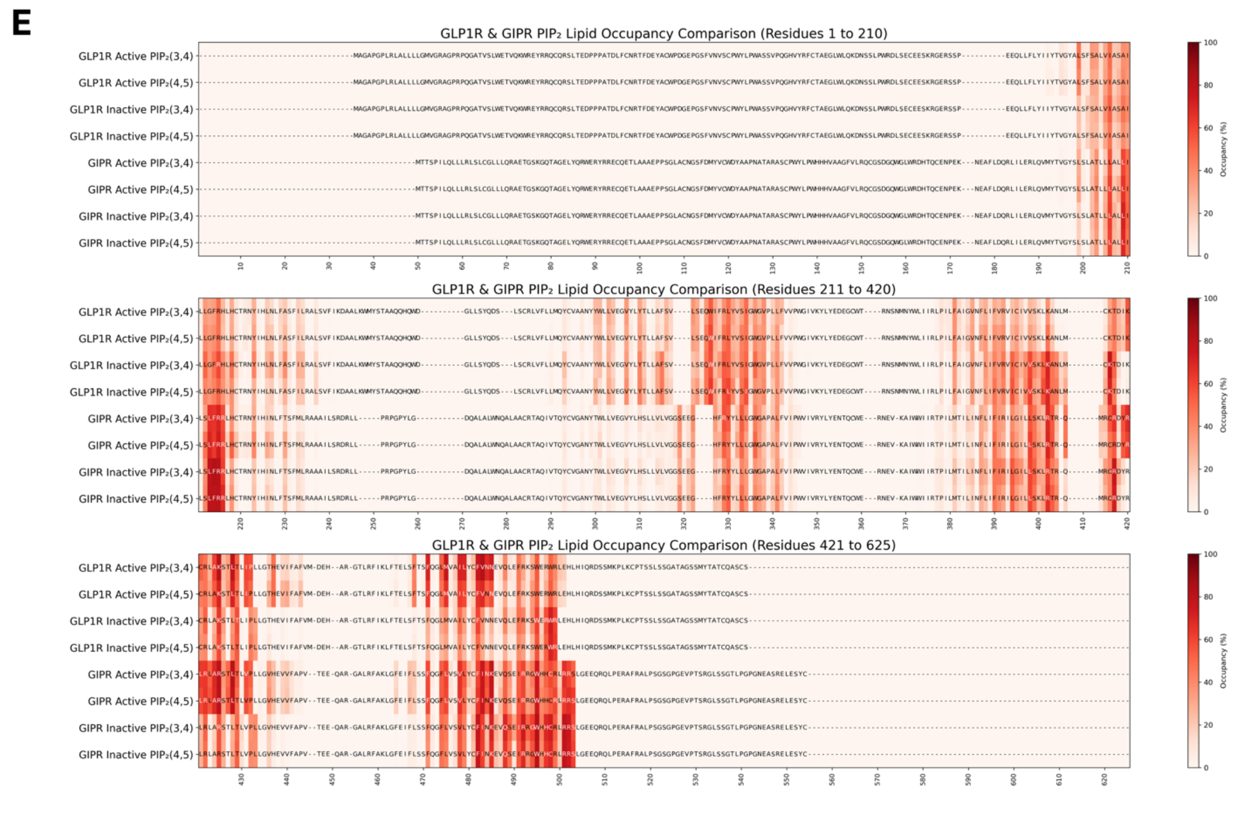


**Fig. S20.**

**Comparison of Martini 3 $PI(3.4)P_2$ and $PI(4,5)P_2$ parameters. A)** Bar chart showing mean $PIP_2$ lipid occupancy per residue for active state GLP1R calculated from three independent simulation repeats (R1, R2, R3) using 7.0 Å cutoff distance for cholesterol. Error bars represent standard deviation across the three repeats. **B**) As for A) but for GLPR1R inactive state. **C**) As for A but for GIPR active state. **D**) As for A) but for GIPR inactive state. **E**) $PIP_2$ occupancy (shown as sequence-based heatmaps (high occupancy to low occupancy: red to white). Cutoff distance was 7 Å. Each heatmap represents averaged data from three simulation repeats.

| | Method | PDB_ID | State | Duration and replicates |
|---|---|---|---|---|
| GLP-1R | cryoEM | 6X18 | Active apo | 10 us x 3 |
| | x-ray | 6LN2 | Inactive apo | 10 us x 3 |
| GCGR | cryoEM | 6LMK | Active apo | 10 us x 3 |
| | x-ray | 5XEZ | Inactive apo | 10 us x 3 |
| GIPR | cryoEM | 7DTY | Active apo | 10 us x 3 |
| | GPCRdb | | Inactive apo | 10 us x 3 |
| GLP-2R | GPCRdb | | Active apo | 10 us x 3 |
| | GPCRdb | | Inactive apo | 10 us x 3 |
| SCTR | cryoEM | 6WZG | Active apo | 10 us x 3 |
| | GPCRdb | | Inactive apo | 10 us x 3 |
| GHRHR | GPCRdb | | Active apo | 10 us x 3 |
| | GPCRdb | | Inactive apo | 10 us x 3 |
| VPAC1 | cryoEM | 8E3Y | Active apo | 10 us x 3 |
| | GPCRdb | | Inactive apo | 10 us x 3 |
| VPAC2 | cryoEM | 7VQX | Active apo | 10 us x 3 |
| | GPCRdb | | Inactive apo | 10 us x 3 |
| PAC1R | GPCRdb | | Active apo | 10 us x 3 |
| | GPCRdb | | Inactive apo | 10 us x 3 |
| PTH1R | GPCRdb | | Active apo | 10 us x 3 |
| | GPCRdb | | Inactive apo | 10 us x 3 |
| PTH2R | GPCRdb | | Active apo | 10 us x 3 |
| | GPCRdb | | Inactive apo | 10 us x 3 |
| CTR | cryoEM | 7TYN | Active apo | 10 us x 3 |
| | GPCRdb | | Inactive apo | 10 us x 3 |
| CLR | GPCRdb | | Active apo | 10 us x 3 |
| | GPCRdb | | Inactive apo | 10 us x 3 |
| CRF1R | GPCRdb | | Active apo | 10 us x 3 |
| | GPCRdb | | Inactive apo | 10 us x 3 |
| CRF2R | GPCRdb | | Active apo | 10 us x 3 |
| | GPCRdb | | Inactive apo | 10 us x 3 |

**Table S1.**
**Models used for each class B1 receptor simulation**.

| GPCR | Repeats | Frame | Membrane composition | Length |
|---|---|---|---|---|
| GLP1R active | R1, R2, R3 | Last frame | 10% GM3 composition[1] | 500 ns |
| GLP1R active | R1, R2, R3 | Last frame | 0% GM3 composition[2] | 500 ns |
| GLP1R inactive | R1, R2, R3 | Last frame | 10% GM3 composition[1] | 500 ns |
| GLP1R inactive | R1, R2, R3 | Last frame | 0% GM3 composition[2] | 500 ns |
| GIPR active | R1, R2, R3 | Last frame | 10% GM3 composition[1] | 500 ns |
| GIPR active | R1, R2, R3 | Last frame | 0% GM3 composition[2] | 500 ns |
| GIPR inactive | R1, R2, R3 | Last frame | 10% GM3 composition[1] | 500 ns |
| GIPR inactive | R1, R2, R3 | Last frame | 0% GM3 composition[2] | 500 ns |
| SCTR active | R1 (frame 1673), R1 (frame 3192), R1 (frame 3421) * | | 10% GM3 composition[1] | 500 ns |
| SCTR inactive | R3 (frame 2460), R3 (frame 3075), R1 (frame 3905) * | | 10% GM3 composition[1] | 500 ns |

1 GM3 composition: The same membrane composition with 10% GM3 as used in the cgMD simulations was used
2 No GM3 composition: The same membrane composition with 0% GM3 (POPC and DOPC were increased to 30%) as used in the cgMD simulations was used
* The selected SCTR frames were the frame (corresponding to time in ns) identified which correspond to the bound pose of the highest residence time site in cgMD.

**Table S2.**
**Details of coarse-grained conversion for atomistic simulations.**
